# Domain motions, dimerization, and membrane interactions of the murine guanylate binding protein 2

**DOI:** 10.1101/2022.04.27.489784

**Authors:** Jennifer Loschwitz, Nora Steffens, Xue Wang, Moritz Schäffler, Klaus Pfeffer, Daniel Degrandi, Birgit Strodel

**Affiliations:** Institute of Theoretical and Computational Chemistry, Heinrich Heine University Düsseldorf, 40225 Düsseldorf, Germany; Institute of Biological Information Processing: Structural Biochemistry (IBI-7), Forschungszentrum Jülich, 52425 Jülich, Germany; Institute of Medical Microbiology and Hospital Hygiene, Heinrich Heine University Düsseldorf, Düsseldorf, Germany

**Keywords:** GBPs, GTPase, MD simulations, GUVs, membrane binding, hinge motion

## Abstract

Guanylate-binding proteins (GBPs) are a group of GTPases that are induced by interferon-γ and are crucial components of cell-autonomous immunity against intracellular pathogens. Here, we examine murine GBP2 (mGBP2), which we have previously shown to be an essential effector protein for the control of *Toxoplasma gondii* replication, with its recruitment through the membrane of the parasitophorous vacuole and its involvement in the destruction of this membrane likely playing a role. The overall aim of our work is to provide a molecular-level understanding of the mutual influences of mGBP2 and the parasitophorous vacuole membrane. To this end, we performed lipid-binding assays which revealed that mGBP2 has a particular affinity for cardiolipin. This observation was confirmed by fluorescence microscopy using giant unilamellar vesicles of different lipid compositions. To obtain an understanding of the protein dynamics and how this is affected by GTP binding, mGBP2 dimerization, and membrane binding, assuming that each of these steps are relevant for the function of the protein, we carried out standard as well as replica exchange molecular dynamics simulations with an accumulated simulation time of more than 30 *μ*s. The main findings from these simulations are that mGBP2 features a large-scale hinge motion in its M/E domain, which is present in each of the studied protein states. When bound to a cardiolipin-containing membrane, this hinge motion is particularly pronounced, leading to an up and down motion of the M/E domain on the membrane, which did not occur on a membrane without cardiolipin. Our prognosis is that this up and down motion has the potential to destroy the membrane following the formation of supramolecular mGBP2 complexes on the membrane surface.

## 1 Introduction

The superfamily of dynamin-like proteins includes the 65–73 kDa guanylate binding proteins (GBPs) that are involved in various structural changes for remodeling cellular membranes or causing fusion/fission events via guanosine-5’-triphosphate (GTP) binding and hydrolysis^1–4^. GBPs are induced by interferon-γ (IFN-γ) and they accumulate at membranes of vacuoles which contain intracellular parasites and bacteria to initiate host cell survival responses^5–9^. Various pathogen-containing vacuoles are targeted by GBPs, including the parasitophorous vacuole membrane (PVM) of the protozoan parasite *Toxoplasma gondii*^10–14^. To date, the process of how the GBPs might bind to and destroy the target’s membrane during the host response as well as the role of GTP binding in it are still poorly understood^10,11,15^. What is clear, however, is that for human guanylate binding proteins GTP binding shifts the monomer/dimer equilibration toward the dimer and even to multimerization^8,16–18^. Until now, seven human GBPs (hGBP1 to hGBP7) and eleven murine GBPs (mGBP1 to mGBP11) are known^19–21^. In this study, we will concentrate on mGBP2, which is the murine orthologue of hGBP1 and was shown to be required to control *T. gondii* replication *in vitro* and in vivo^11^. In order to execute their antimicrobial function, murine GBPs can specifically associate with pathogen containing compartments of intracellular microbes, which encompass pathogen-containing vacuoles as well as viral replication compartments^11,12,22–25^, resulting in the innate immune detection and cell-autonomous clearing or of vacuole-resident pathogens^7,24,26,27^. After IFN-γinduction, mGBP2 molecules assemble in vesicle-like structures in the cytoplasm and, after *T. gondii* entry into the host cell, mGBP2 attacks the PVM as orchestrated, supramolecular complexes, ultimately promoting the destruction of the vacuolar membrane^7,11^. This stage is followed by relocation of mGBP2 to the plasma membrane of the intracellular parasite, which in turn likely leads to its destruction^7^. Importantly, inactivation of mGBP1, mGBP2, or mGBP7 in mice has been shown to result in a vastly impaired resistance to infections with type II *T. gondii*^11,24^.

The structure of mGBP2 was not determined yet by experimental means, but can be modeled using its homology to hGBP1 with a sequence identity of 66.2% for the whole protein. The structure of hGBP1 has been resolved as nucleotide-free (PDB ID 1DG3, 592 residues)^28^ and nucleotide-bound monomer (PDB ID: 1F5N)^29^. Generally, GBPs have between 586 and 638 residues, and contain three domains, which is illustrated for mGBP2 in Fig. 1**a**. The proteins have a GTPase (G), a middle (M) and an effector (E) domain. The N-terminal GTPase domain is generally large (residue 1–303 in mGBP2), binds GTP and hydrolyzes it to GMP via GDP as intermediate. The middle domain (residues 304–481) gives the elongated structure and finally, the effector domain (482–589) is responsible for membrane interaction.

**Figure 1.**
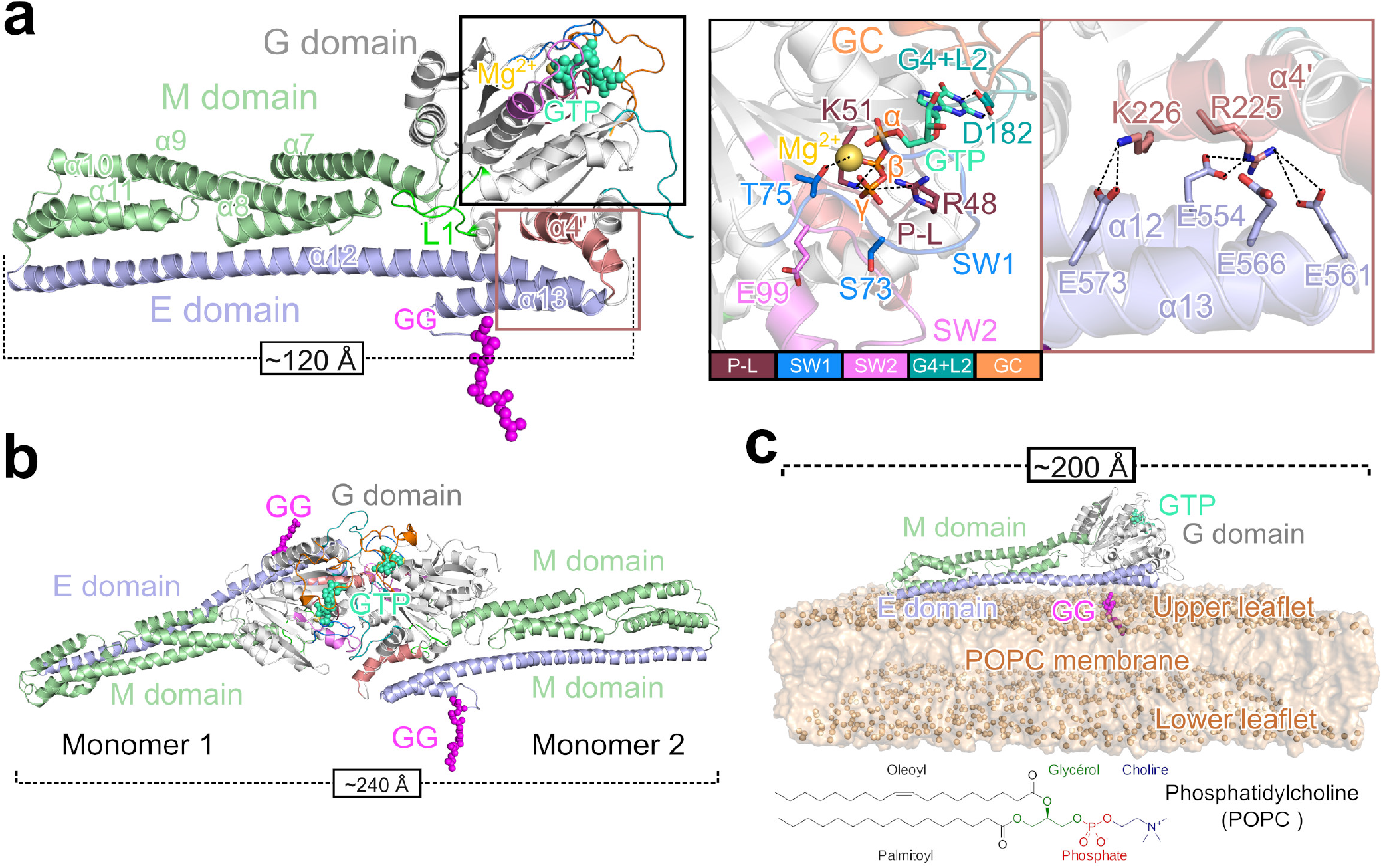
Model systems of mGBP2 investigated in this study. **(a)** The homology model of the entire mGBP2 molecule contains three different domains: (i) the G domain (gray cartoon) with GTP (green spheres) and Mg^2+^ (orange sphere); (ii) the M domain (green cartoon); (iii) the E domain (blue cartoon), and the geranylgeranyl lipid anchor (GG, magenta spheres). The black frame shows a zoomed view of the GTP binding site with the different G motifs and loops being highlighted (see color code at the bottom). Moreover, the residues important for GTP binding and hydrolysis are indicated too, along with their interaction (dashed black lines) with the different parts of GTP (*α*-, *β* - and *γ*-phosphate) or with Mg^2+^. The brown frame shows the interactions between the *α*4’ of the G domain and *α*12/13 of the E domain. The side chains of the important residues are shown as sticks and salt bridges are indicated by dashed black lines. **(b)** The model of the mGBP2 dimer, that includes both GTP and the geranylgeranyl group, is based on the crystal structure of the G domain dimer of hGBP1 (PDB ID: 2BC9)^31^. **(c)** The membrane-associated mGBP2 monomer model is anchored via the geranylgeranyl group to a POPC membrane (1-palmitoyl-2-oleoyl-sn-glycero-3-phosphocholine, brown surface, phosphor atoms highlighted as spheres). The chemical structure of POPC is given at the bottom.

The GTP binding site in the G domain involves four conserved sequence elements called G motifs (Fig. 1**a**, Supplementary Table S1). These are: the canonical G1 motif GXXXXGK(S/T), also called phosphate-binding loop (P-L), the switch1/G2 (SW1) motif, the phosphate- and Mg^2+^-binding switch2/G3 (SW2) DXXG sequence, and the nucleotide-specificity providing G4 motif X(V/L)RD with the loop 2 (G4+L2^)10,21,30^. The phosphate-binding loop has an integrated arginine finger R48 and the residue K51 for GTP-hydrolysis, where the K51A mutant leads to a dysfunctional mGBP2^10^. Like hGBP1 and also mGBP7, mGBP2 can hydrolyze GTP to GMP via GDP by moving the *β*-phosphate of GDP into the γ-phosphate position for the hydrolysis^10,30–32^. This contrasts with other GTPases, such as Ras and other dynamin-like proteins, where the hydrolysis is limited to GDP. Simulations of hGBP1 showed that S73 and T75, which are part of the SW1 motif, form a hydrogen bond network together with E99 in SW2 and water molecules, which stabilizes the Mg^2+^ ion affiliated with the γ-phosphate of GTP^32^. It was further found that while E99 and S73 are not directly involved in the hydrolysis, replacing them with alanine reduces the enzymatic efficiency^33^, which was also observed in experiments of mGBP2^10^. Finally, D182, which is part of the G4+L2 motif shall be specially mentioned, as it is responsible for the nucleotide preference for GTP in mGBP2^10^ and in hGBP1^29^. Another key structural element of the G domain is the guanine cap (GC) involving residues 235–256. From the crystal structure of the G domain dimer of hGBP1 (PDB ID: 2BC9) it is known that the protein dimerization is mainly realized by the two guanine caps interacting with each other, involving a transition toward a closed conformation of this loop region^31^

The M domain of both hGBP1 and mGBP2 is a two-helix bundle (*α*7, *α*8 and *α*10, *α*11) connected with a long helix (*α*9). This bundle of helices gives the GBPs an elongated shape, at up to 120 Å long, and stabilizes the *α*12 of the E domain (Fig. 1**a**, Supplementary Table S1)^28^. The adjacent E domain with the long helix *α*12 and a short helix *α*13 is stretched back to the G domain along to the M domain, which is stabilized by contacts with both the M and the G domain. Of particular interest are salt bridges formed between *α*4’ of the G domain and *α*13 as well as hydrophobic contacts between loop 1 of the G domain and the E domain (Fig. 1**a**). In mGBP2 these electrostatic contacts involve residues R225 and K226 of *α*4’, E554 and E561 of *α*12, and E566 and E573 of *α*13. In hGBP1, the analogous interactions likely break during the conformational change following GTP binding^8,34,35^.

In uninfected, interferon-*γ* stimulated cells, mGBP2 localizes in vesicle-like structures within the cytosol. Moreover, imaging pictures revealed an accumulation of mGBP2 at the PVM of *T. gondii*^7,10,36^, indicating the need to also study the multimerized state of mGBP2. It was already demonstrated that, like hGBP1, mGBP2 forms dimers and larger multimers in a GTP-dependent manner^7^ and that dimerization has a strong effect on the self-activation of GTP hydrolysis, in which two arginine residues that are part of the dimer interface, R240 and R244 play a key role^37^. However, the only high-resolution structural information that is available for GBP multimers is the crystal structure of the G domain dimer of the hGBP1 (PDB ID: 2B92)^31^. Based on this structure we built models for the full-length dimers of hGBP1^35^ and mGBP2 (Fig. 1**b**), which span a length of about ~240 Å. The dimerization interface mainly involves the guanine cap regions of both proteins. Association with the membrane of the parasitophorous vacuole is most likely mediated by the C-terminal lipid anchor that mGBP2 features, as does hGBP1. Both proteins have a CaaX motif for post-translational modification by a farnesyl group (hGBP1) or a geranylgeranyl group (mGBP2)^11,19,34,38,39^. This anchor lies at the end of the E domain; in the case of mGBP2 it is attached to C586 (Fig. 1**c**).

The aim of this study is to analyze the conformational dynamics of mGBP2 and how the protein motions are affected by GTP binding, post-translational modification, membrane binding, and dimerization. To this end, we performed all-atom molecular dynamics (MD) simulations, employing the Hamiltonian replica exchange molecular dynamics (HREMD) technique to enhance the conformational sampling. To mimic the different protein states, we simulated the apo-protein that has neither GTP bound nor the lipid anchor attached (mGBP2_apo_), the protein with GTP being bound (mGBP2_GTP_), and the protein that has GTP bound and the geranylgeranyl anchor attached, which we denote as the holo-state (mGBP2_holo_). As mGBP2 is the orthologue of hGBP1, for which we previously uncovered a large-scale hinge motion involving the M and E domains^35^, we paid particular attention to identifying similar motions in mGBP2 and how they are affected by GTP binding and the presence of the geranylgeranyl anchor, as this has not been studied before. We further study the binding of mGBP2 to lipid membranes. Since the lipid composition of the PVM is not known, we first used membrane lipid strips to uncover the preferred lipid binding partner of mGBP2, which turns out to be cardiolipin. This finding we then confirmed using confocal microscopy applied to mGBP2 tagged with green fluorescent protein (GFP) in the presence of giant unilamellar vesicles (GUVs). To understand the effects of membrane binding to the conformational dynamics of mGBP2 and the role that cardiolipin plays in it, we simulated the protein anchored to a simple 1-palmitoyl-2-oleoyl-sn-glycero-3-phosphocholine (POPC) membrane and a mixed membrane containing cardiolipin. Finally, the dynamics of the mGBP2 dimer is explored. The simulations of these different systems give us fundamental insights into the dynamics of mGBP2, which is of relevance for its biological functions.

## 2 Results

### 2.1 Monomeric mGBP2 in solution

We started the simulations with the monomeric mGBP2 in its apo-state in solution. This serves as a reference to delineate the effects of GTP binding, geranylgeranylation, protein dimerization, and membrane binding by successively including these aspects in the simulations following thereafter. Because mGBP2 is a large protein with almost 600 residues, we performed enhanced sampling by running Hamilitonian replica exchange molecular dynamics (HREMD) simulations involving 16 replicas with 200 ns per replicate simulation (amounting to 3.2 *μ*s cumulated simulation time) for each of the monomer systems to better sample their conformational space. For the analysis, we use the target replica simulated with an unmodified energy function. First, we unravel the overall motions and then investigate in more detail how the different regions of mGBP2 are affected by GTP binding and geranylgeranylation.

#### 2.1.1 Overall flexibility

To evaluate the protein flexibility, we calculated the root mean square fluctuations (RMSF) of the C_α_ atoms. Fig. 2 shows that GTP and geranylgeranyl have only minor effects on the overall dynamics of mGBP2. For being better able to correlate the dynamics with the mGBP2 structure, we projected the RMSF values onto the structure from which the MD simulations were started (Supplementary Fig. S1**a–c**). We observed that mGBP2 exhibits very similar dynamics to hGBP1,^35^ with the core of the G domain being the least flexible. The most flexible part is the tip of the M and E domain (around residue 480) and the neighboring helices *α*10 (residues 420–446) and *α*12 (residues 482–561). The dynamics of the M and E domain (M/E domain) derives from a hinge motion of both domains, which will be discussed below. However, the M/E domain of mGBP2 is stiffer than that of hGBP1, as the latter reached RMSF values of up to 16 Å, compared to only 6 Å in mGBP2.

**Figure 2.**
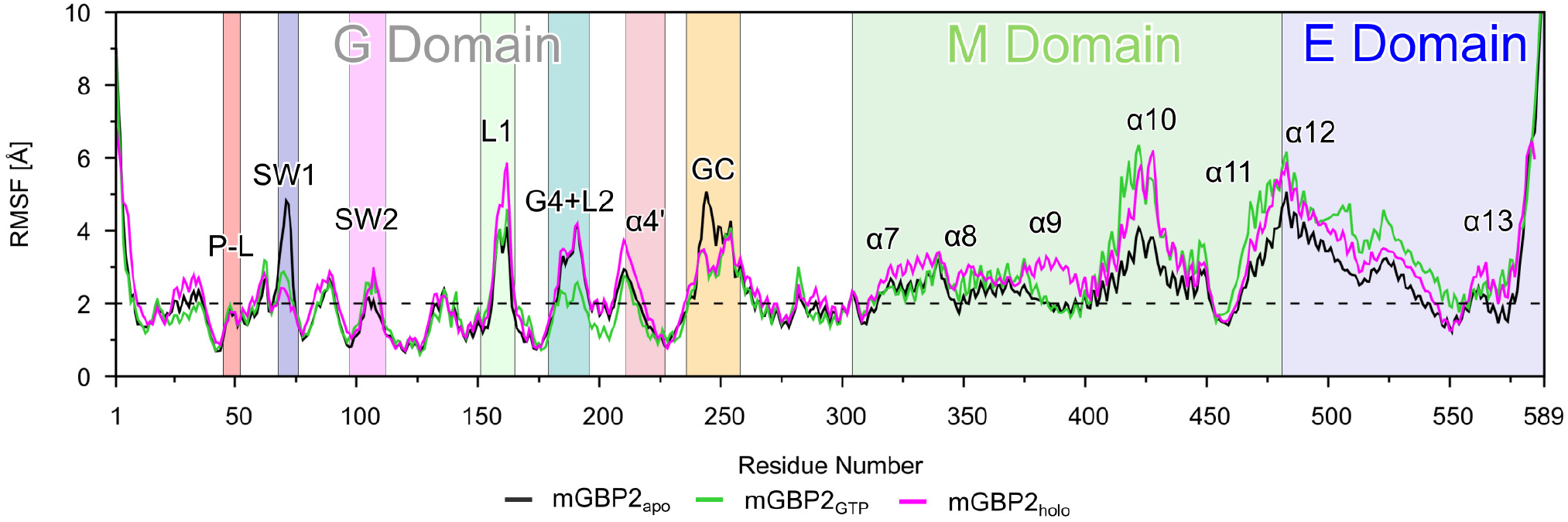
The root mean square fluctuations per residue of monomeric mGBP2 obtained from HREMD simulations of mGBP2_apo_ (black), mGBP2_GTP_ (green), and mGBP2_holo_ (magenta), as sampled in the target replica. All important motifs, loops, and helices are labeled and a background color added using the same colors as for the corresponding structural units as in Fig. 1.

#### 2.1.2 Effects of GTP and geranylgeranyl binding on the G domain

The dynamics of the G domain loops in mGBP2 without GTP are very similar to their dynamics in the GTP-free hGBP1^35^. In particular, the SW1 and guanine cap are very flexible with RMSF values above 5 Å, while GTP binding makes these loops more rigid (Fig. 2). To quantify the dynamics of the G domain, we clustered the conformations of each loop by using a cutoff of 2.5 Å with a fitted trajectory on the β-sheets of the G domain. The results of this analysis are summarized in Table 1 and shown in Fig. 3.

**Figure 3.**
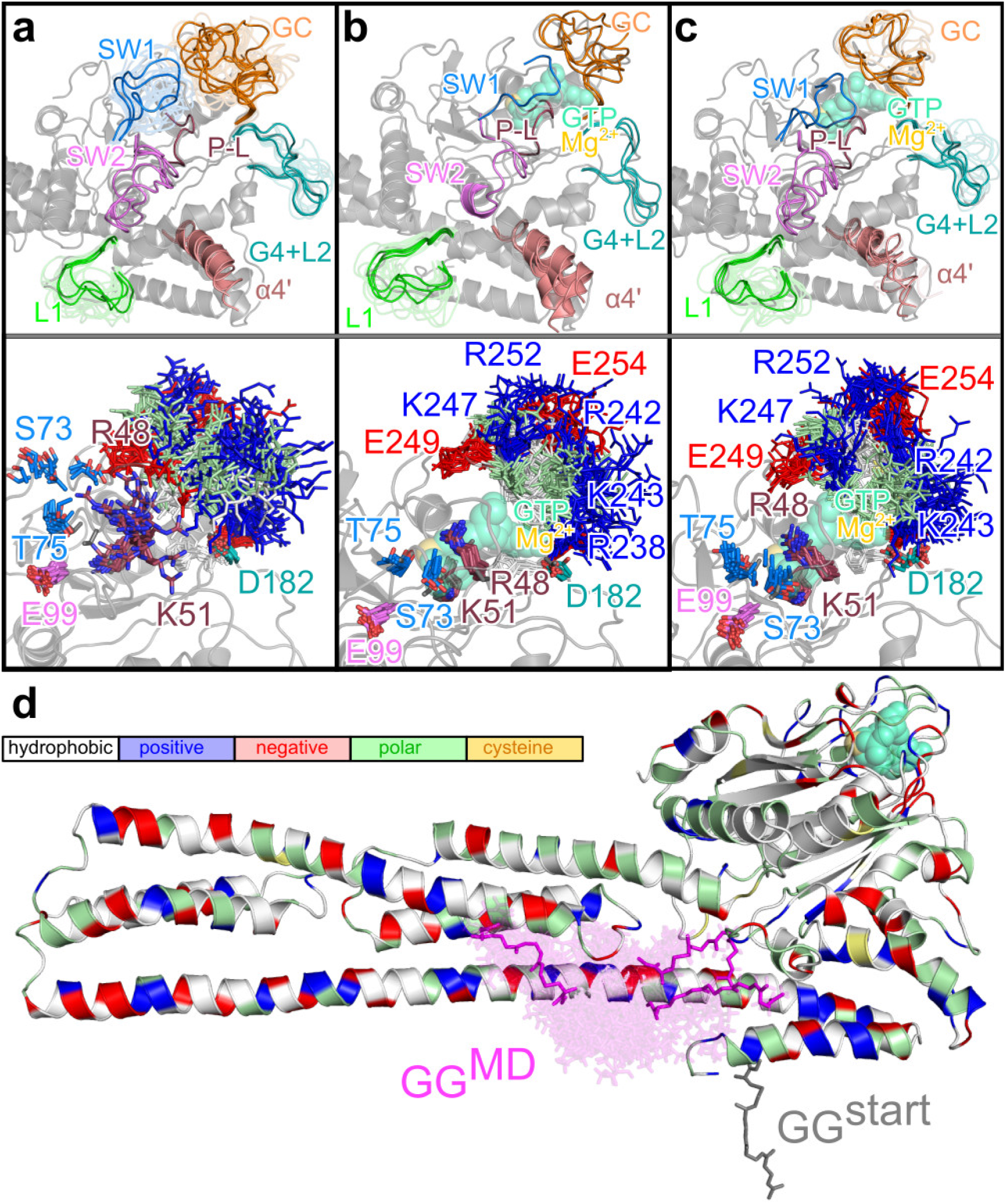
Conformational clusters of the G domain loops of (a) mGBP2_apo_, (b) mGBP2_GTP_, and (c) mGBP2_holo_ as well as (d) for the geranylgeranyl anchor. **(a-c)** In the upper panels, the central conformations for the five most populated clusters (or fewer if < 5 were found) are shown in opaque and for the other clusters in transparent, using different colors for the different loops as specified in the figures. For mGBP2_GTP_ and mGBP2_holo_, the GTP-binding pocket is indicated by showing GTP and Mg^2+^ (in transparent green and light orange). The helix *α*4’ is also highlighted in light red. The homology model of mGBP2 is shown as reference in a gray cartoon representation. In the lower panels, the different conformations of key residues for all clusters are shown, as sticks colored according to their residue type (white: apolar; green: polar; blue: positively charged; red: negatively charged), and labeled. **(d)** The central cluster conformations of the geranylgeranyl group (GG, magenta sticks) of mGBP2_holo_ are shown in opaque for the three most populated clusters and in transparent for all other clusters. The initial conformation of the geranylgeranyl group used to start the HREMD simulation is also depicted (gray sticks), while the homology model of the whole protein is presented as cartoon and colored based on residue type as explained in the legend box.

**Table 1.**
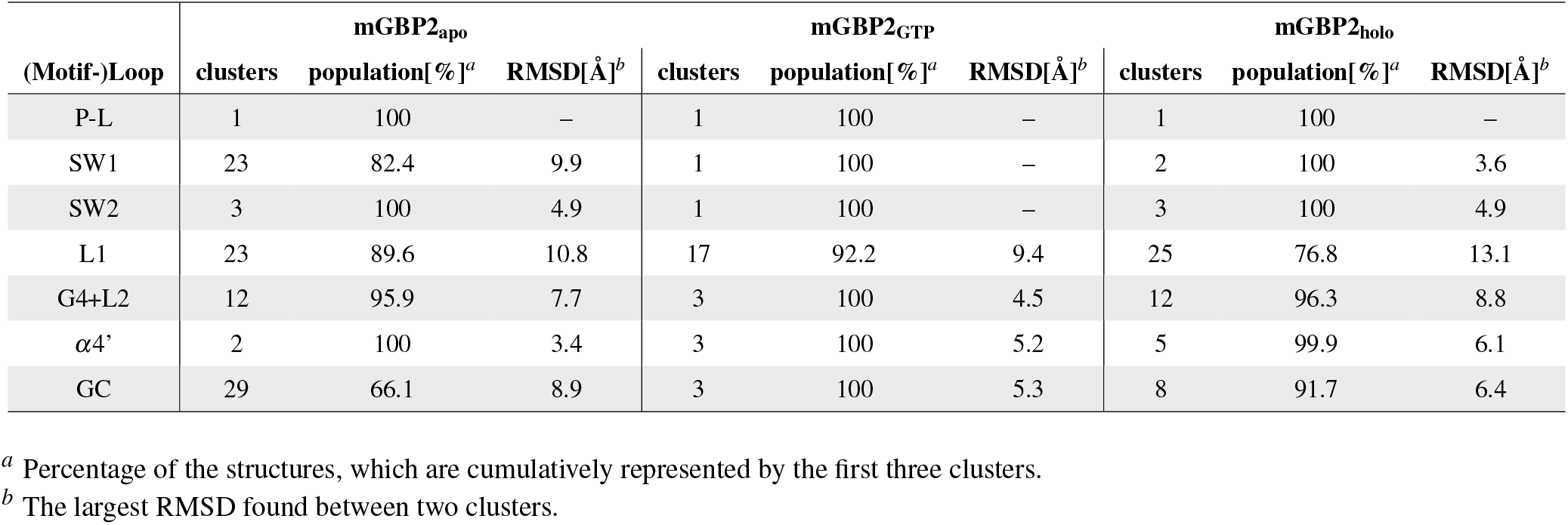
Flexibility of the G domain loops of monomeric mGBP2 determined by clustering of the conformations sampled in the HREMD target replica.

The two most flexible loops are SW1 and the guanine cap, which can both switch between open and closed forms and adopt all kinds of intermediate states. In SW1, residue S73 that is important for GTP binding by stabilization of Mg^2+^ is flexible in the absence thereof and can also interact with R48, which is the arginine finger needed for the GTP hydrolysis. Without GTP being bound, R48 is very flexible too and adopts conformations covering the whole GTP binding site, thereby also interacting with the negatively charged residue E249 of the guanine cap and E105 of SW2. The flexibility of SW1 creates space for the the guanine cap in the GTP binding site, which can therefore adopt all kinds of conformations (Fig. 3**a**). In contrast, the residues K51 in the phosphate-binding loop, T75 in SW1, E99 in SW2, and D182 in the G4 motif/L2 loop are not very flexible despite the absence of GTP. These residues were also fixed in the GTP-free hGBP1, from which we concluded that the phosphate-binding loop and SW2 do not require GTP for adopting a closed conformation as they can be closed without GTP^35^. However, GTP binding is assumed to shift the equilibrium toward the closed states of SW1 and the guanine cap.

The current results obtained for mGBP2 in the presence of GTP confirm that this is indeed the case. In mGBP2_GTP_, SW1 adopts only a single conformation, which is the closed one that is stabilized by key interactions: R48, S73 and K51 interact with the γ-phosphate of GTP and T75 has contacts with Mg^2+^ (Fig. 3**b**). Residue D182 (in the G4 motif/L2 loop), on the other hand, switches between conformations, where it interacts with GTP, and orientations that are turned away from GTP for interacting with R238 in the guanine cap. Residue E99 (in SW2) is always pointed away from GTP. For hGBP1 it was shown that E99 instead forms a hydrogen-bond network involving two water molecules and S73^32^.

As a result of the stable interactions and conformations of S73 and R48, they are no longer available for interactions with residues of the guanine cap. This, in turn, causes the guanine cap to adopt an ordered conformation, which is stabilized by interactions among the guanine cap residues, in particular between the the positively charged residues R242, K243, K247, and R252 that form salt bridges with the negatively charged residues E249, E253, E254, E255 (Fig. 3**b**). When the geranylgeranyl group is added to the picture, leading to mGBP2_holo_, most loops are slightly more flexible than those in mGBP2_GTP_ (Fig. 3**c**). We will see shortly that this is because of interactions of the geranylgeranyl lipid anchor with the G domain. Nonetheless, despite this increased flexibility, the two conformational clusters identified for SW1 in mGBP2_holo_ all belong to the closed state. The phosphate-binding loop containing the residues R48 and K51 for GTP hydrolysis is already quite stable without GTP being bound. In mGBP2_GTP_ and mGBP2_holo_ this loop becomes even more rigid (RMSF values under 2 Å). In both cases, only one conformational cluster was found for the loop. Similarly, SW2 and D182 from the G4 motif also rigidified in the presence of GTP. Thus, the interplay between the G domain and GTP as well as Mg^2+^ stabilize the loops surrounding the GTP binding site in a conformation optimal for the subsequent hydrolysis reaction. Different behavior between mGBP2_GTP_ and mGBP2_holo_ is observed for the G4 motif/L2 loop. Only in the case of mGBP2_GTP_, does GTP binding cause this loop region to become stable; only three conformational clusters are being found here (Table 1). This stability results from a β-hairpin conformation that is formed in L2, which in turn includes three stable backbone hydrogen bonds between L187 and V194, V189 and K192, and S196 and L200 as well as a hydrogen bond between the side chain or backbone of T195 and the side chain of E198. This conformation is further stabilized by a hydrophobic cavity made of the residues of a short helix and the adjacent β-hairpin, consisting of F183, L187, V189, V194, Y199, and I235. In mGBP2_holo_ and mGBP2_apo_, on the other hand, this G4 motif/L2 loop is flexible with RMSF values between 3 and 4 Å leading to 12 conformational clusters in either case. Here, the β-hairpin is only intermittently adopted.

Next, we turn our attention to *α*4’, as this helix was suggested to undergo conformational changes upon GTP binding and hydrolysis^34^. However, even in GTP-free mGBP2 we find *α*4’ to be flexible. In fact, in each system, this helix prefers a position slightly different from the one predicted by homology modeling (Fig. 3). In the presence of GTP and the geranylgeranyl group, i.e., in mGBP2_holo_, *α*4’ is particularly flexible, which is visible from its RMSF values of up to 4 Å (Fig. 2). Moreover, it can move up to 18 Å (Supplementary Fig. S2**c**), causing its reversible unfolding between residues K212–Q219. This instability of *α*4’ results from the geranylgeranyl group interacting with loop L1, which in turn affects *α*4’. In mGBP2_holo_, loop L1 is the most flexible, as the number of loop L1 clusters is 25 in mGBP2_holo_, 23 in mGBP2_apo_, and 17 in mGBP2_GTP_. Moreover in mGBP2_holo_, the first three clusters represent only 77% of the L1 conformations, while over 95% of the structures are included in the first three L1 clusters in mGBP2_apo_ and mGBP2_GTP_.

Finally, we assess the effect of GTP on the guanine cap, which we identify as not as strong as for the other motifs/loops that are in direct contact with GTP. However, as already explained above, as a consequence of the other motifs/loops becoming more ordered upon GTP binding, this inflicts a certain degree of order in the guanine cap too. Fig.**3b/c** shows that in mGBP2_GTP_ and mGBP2_holo_ the guanine cap adopts a conformation that is closing off the GTP binding site. The clustering analysis also reveals that this loop also becomes more rigid, as evidenced by the number of conformational clusters decreasing from 29 in mGBP2_apo_ to 3 and 8 in mGBP2_GTP_ and mGBP2_holo_, respectively. The increased stability of the guanine cap following GTP binding should facilitate the dimerization of mGBP2, as the protein-protein interface in the dimer mainly involves the guanine caps of both proteins. Because GTP binding shifts the conformational equilibrium of the guanine cap already in the monomer towards the one in the dimer, it can be expected that this conformation shift facilitates protein-protein recognition for dimer formation. Indeed we do demonstrate this to be the case in the dimer analysis section below.

In summary, all loops of the G domain become more rigid upon GTP binding. Some of the conformational stiffening is being undone by geranylgeranylation, which especially applies to loops L1 and G4+L2 as well as helix *α*4’, due to their direct or indirect interactions with the geranylgeranyl group. Overall, mGBP2_GTP_ seems the most stable monomeric protein state compared to the two other states, mGBP2_apo_ and mGBP2_holo_.

#### 2.1.3 Motions of the geranylgeranyl lipid anchor

To better understand the effects of the geranylgeranyl lipid anchor on the structural dynamics of mGBP2, we analyze its motions in detail. We attached the geranylgeranyl group at the C-terminal end of mGBP2 such that it pointed away from the protein (Fig. 1**a**). However, from this position it readily moved away, during the HREMD simulation of mGBP2_holo_, in order to avoid being completely water-exposed as it has mostly hydrophobic moieties. The clustering analysis revealed the geranylgeranyl anchor to prefer positions close to helices *α*7, *α*9 and *α*12, whose hydrophobic residues attract the anchor (Fig. 3**d**). Interestingly, the geranylgeranyl anchor did not move towards the most hydrophobic part of mGBP2, which is a hydrophobic cave formed by the G domain helices *α*4’ and *α*6 as well as L1 and the *β*-sheet behind them, along with the first half of *α*7 and the loop leading to *α*7. Nonetheless, the geranylgeranyl anchor does interact with the L1 loop, which, as a result, becomes more flexible (as discussed above). The interactions of the geranylgeranyl anchor with the *α*7, and also the helices *α*9, *α*10, and *α*13, which are all part of the M or E domain, cause an increase in the motions of these two domains, which will be analyzed in more detail below. The geranylgeranyl anchor is rather flexible, as supported by 79 conformational clusters representing only 39.4% of the geranylgeranyl conformations that we identified from applying conformational clustering with a 2.0 Å cutoff to the HREMD simulation of mGBP2_holo_ (Table 1 and Fig. 3**d**). This flexibility derives from the conformational freedom of the last seven residues (K578–K585) of mGBP2 that are located between α 13 and the geranylgeranyl anchor.

Taken together, these data indicate that the geranylgeranyl anchor is highly flexible and moves to a more hydrophobic cavity formed by residues of the M and E domain. However, it does not constantly bind to specific residues but it keeps its flexibility.

#### 2.1.4 Effects of GTP binding and geranylgeranylation on the M/E domain

Next, we had a closer look at how GTP and the geranylgeranyl lipid anchor influence the M/E domain of mGBP2. Therefore, we applied principal component analysis (PCA) to the M/E domain motions. In the following we focus on the two eigenvectors PC1 and PC2 that represent the domain’s two main motions. In order to allow for a direct comparability between the eigenvectors of the three monomeric systems under study, i.e., mGBP2_apo_, mGBP2_GTP_, and mGBP2_holo_, we applied the PCA to the concatenated target-replica trajectories of the three HREMD simulations and then projected the individual trajectories onto the common eigenvectors. The resulting plots are shown in Fig.**4a** and characteristics of these projections are summarized in Supplementary Table S2. In general, the distributions of the conformations along PC1 and PC2 are very similar in the three systems. The main difference between them is the amplitude of the motions. In the mGBP2_apo_, the value range is between –12.1 nm^2^ (PC1_min_) and 8.5 nm^2^ (PC1_max_) for PC1 as well as –8.1 nm^2^ (PC2_min_) and 10.9 nm^2^ (PC2_max_) for PC2, which serve as reference. Upon GTP binding, the deflection along PC1 increased, with PC1_max_ reaching 12.7 nm^2^ (ΔPC1_max_ = 4.2 nm^2^), whereas all other values lie in the same area as for mGBP2_apo_. When the geranylgeranyl anchor is further added to the protein, this main motion became even more pronounced: the mGBP2_holo_ has an almost two-fold higher PC1_max_ value of 16.9 nm^2^ (ΔPC1_max_ = 8.4 nm^2^) compared to mGBP2_apo_. On the other hand, the motion along PC2 is somewhat reduced in this protein, as PC2_max_ only reaches a value of 9.4 nm^2^ (ΔPC2_max_ = –1.5 nm^2^). In order to depict the two main motions of the proteins, in Fig.**4b** the mGBP2 conformations corresponding to the minimal and maximal PC1/PC2 values are shown. One can see that the motion along PC1 stems from a hinge movement of the M/E domain, similar to the one that we had already identified for hGBP1^35^. The motion along PC2, on the other hand, is a twist movement of the M/E domain. Hence, addition of GTP and geranylgeranylation have a particular enhancing effect on the hinge motion. This is not only confirmed by the larger RMSF values observed for the M/E domain of the mGBP2_GTP_ and mGBP2_holo_ compared to the mGBP2_apo_ (Supplementary Fig. S1**a–c**), but are also witnessed by the maximal deviations of the tip of the M/E domain from the starting structure of the HREMD simulations (~ 33 Å for mGBP2_apo_ vs. > 42 Å for mGBP2_GTP_ and mGBP2_holo_).

**Figure 4.**
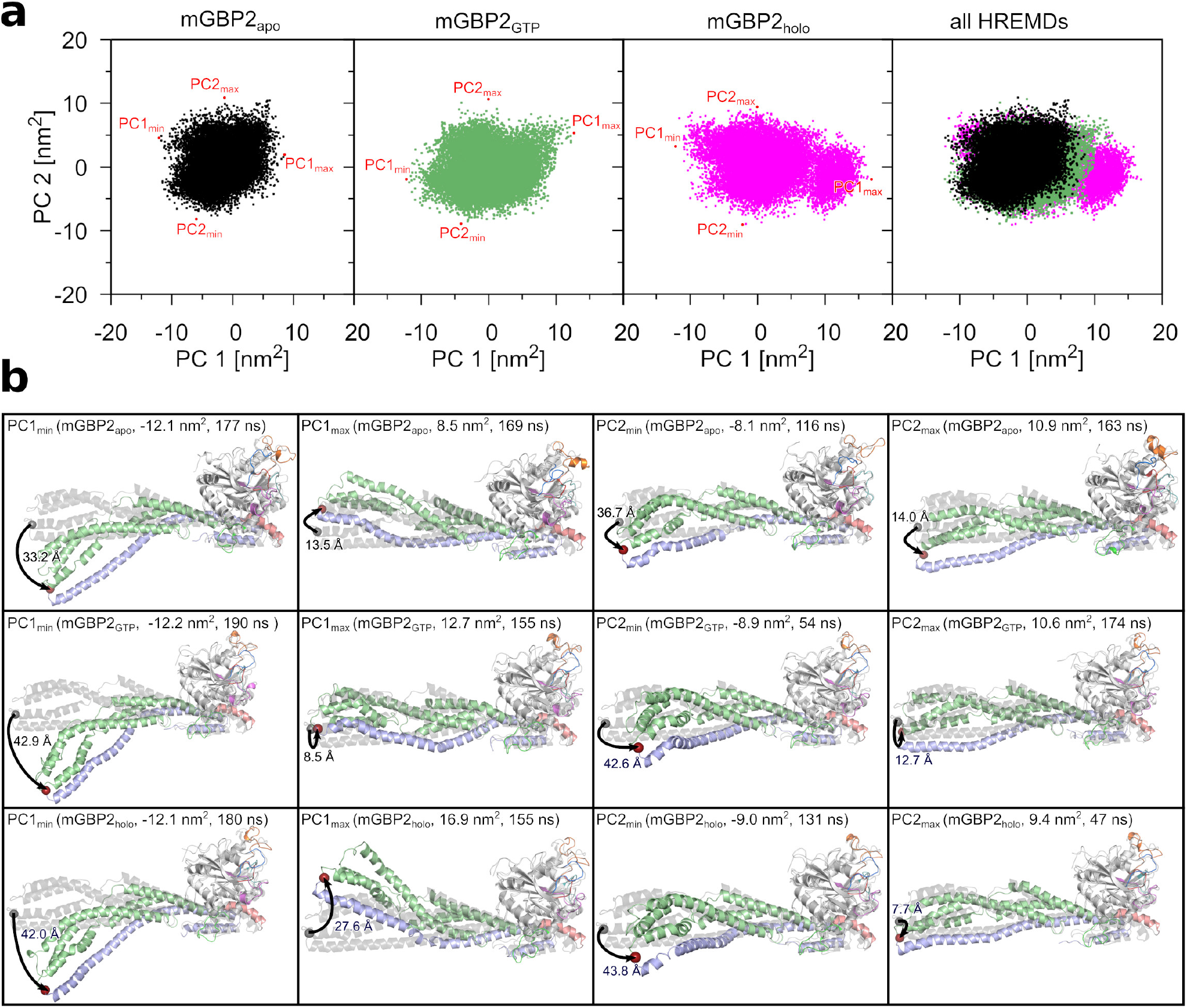
Main motions of the M and E domain in monomeric mGBP2 identified by PCA of the target replica of the HREMD simulations. **(a)** The projection of the conformations collected in the target replica onto the first two principal components (PC1 and PC2). The results for GTP-free mGBP2 (black), mGBP2 with GTP bound (green), and mGBP2 with both GTP and geranylgeranyl anchor (magenta) are shown from left to right. For a better comparability, all three projections are shown together in the rightmost panel. The minimum and maximum values of the projections are indicated by red dots and labeled. **(b)** The conformations at the minimal and maximal PC1 and PC2 values are shown for the mGBP2_apo_ (top), mGBP2_GTP_ (middle) and mGBP2_holo_ (bottom). The corresponding PC1 and PC2 values are provided too, along with the simulation time at which these conformations were sampled. The motions of the tip of the M/E domain relative to the starting structure (gray cartoon) are indicated by arrows and the maximal amplitudes (in Å) are given.

To further quantify the motions of the M/E domain, we calculated the change in Cartesian coordinates (Δ*x*, Δ*y*, Δ*z*) of residue L480, which defines the tip of the M/E domain. Fig.**5a** shows the reference conformation in the Cartesian room, with the location of the L480 being highlighted. Fig.**5b** reveals that the hinge motion is mainly characterized by motions into positive y direction, corresponding of the tip of the M/E domain moving toward the G domain. The average values for the mGBP2_apo_ are Δ*x* = −4.2 ± 0.02 Å, Δ*y* = 26.0 ± 0.07 Å, and Δ*z* = −6.4 ± 0.02 Å. In Supplementary Table S2, also the minimum and maximum changes are listed. For the mGBP2_GTP_, the values are in a similar range, however, the motion into negative Δ*y* direction has grown. Here, values down to −19.0 Å are reached, compared to −8.1 Å in mGBP2_apo_, which means that the motion can also go into the opposite direction with the tip of the M/E domain moving away from the G domain side that interacts with the *α*13. With the hinge motion taking place into the opposite direction, the larger PC1 space as seen for mGBP2_GTP_ compared to mGBP2_apo_ can be explained. Moreover, in mGBP2_GTP_ more distinct motions along the z direction are observed too. In the presence of the geranylgeranyl anchor, these two effects are further increased, especially Δ*z* values are even larger. The most extreme motions of the M/E domain are shown by representative snapshots in Supplementary Fig. S2. They show that in the presence of GTP and the geranylgeranyl anchor, the helices *α*9 and *α*12 can (reversibly) unfold, involving residues R491–E503 (Supplementary Figs. S1**a–c** and S2**b/c**), leading to a drop in the average helix content in the E domain by 3–4% relative to 88% in mGBP2_apo_ (Supplementary Table S2). Hence, GTP binding and the addition of the geranylgeranyl anchor have an influence on the motions of the M/E domain. In the case of the geranylgeranyl anchor, these effects emerge from the direct interactions between that anchor and the M/E domain, whereas in the case of the GTP binding it could be an allosteric effect that couples the G domain with the M/E domain.

**Figure 5.**
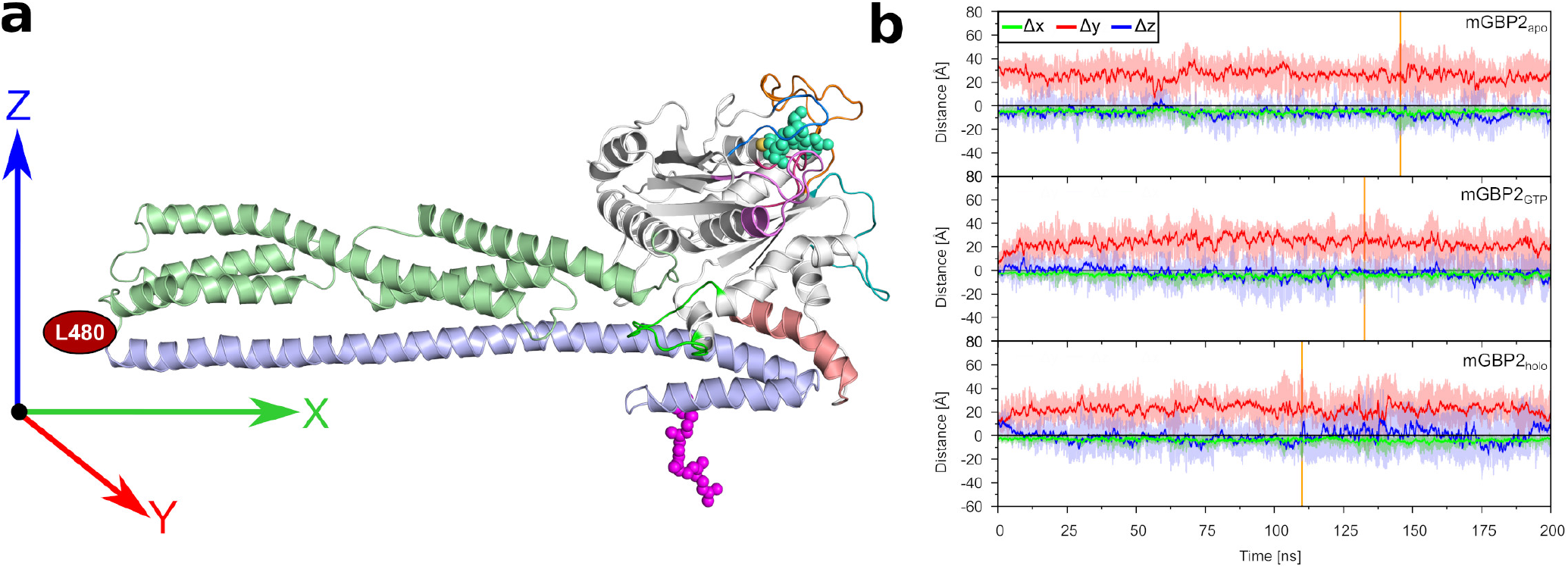
Motions of residue L480 in the HREMD simulations of monomeric mGBP2. **(a)** Reference conformation of the mGBP2 is shown for the calculation of the coordinate distances difference (Δ*x*, Δ*y*, Δ*z*) of L480. **(b)** The evolution of these distances (Δ*x*: green, Δ*y*: red, Δ*z*: blue) is shown for mGBP2_apo_ (top), mGBP2_GTP_ (middle), and mGBP2_holo_ (bottom). The solid lines show the running average and the shades illustrate the raw data. Furthermore, the zero line is shown in black, whereas the vertical orange line indicates the frame with the maximal distance of the L480 residue between reference structure and 200 ns HREMD simulations.

To examine the possibility of an allosteric effect, we calculated transition networks (TNs) using our software ATRANET^40–43^ to reveal conformational transition pathways within mGBP2. Since HREMD data does not allow the construction of TNs due to the exchanges between replicas causing the loss of the information about time correlation between states, we run standard MD simulations of 1 *μ*s length for both mGBP2_apo_ and mGBP2_holo_. A TN needs molecular features (or descriptors) as input and we tested different descriptor combinations. The most informative TNs were obtained for following three descriptors: i) the state of the three possible salt bridges R225–E554, R225–E561, and K226–E573 which link *α*4’ with *α*12/13 and might allow for allosteric communication between the G and the E domain, ii) the RMSD of the guanine cap (RMSD_GC_) as a reporter for GTP binding, and iii) the motion of the tip of the M/E domain as measured be the distance of residue 480 with respect to the homology model that was used for starting the MD simulations (d480). The resulting TNs are shown in Supplementary Fig. S3. For further analysis, we identified local communities of highly interconnected states, which we call macrostates here. We found three macrostates for mGBP2_apo_ and four for mGBP2_holo_.

One main difference between the two TNs is the vertical spread of states, which mainly separates the states based on the RMSDGC. Without GTP, RMSDGC ranges from 0 to 8 Å, whereas for mGBP2_holo_ it only goes up to 5 Å resulting in a narrower spread of the states along the vertical. The range of values for d480 is similar in both networks, which can be as high as 46 Å, but is on average lower in mGBP2_holo_. This finding, which at first might seem surprising considering that the PCA and Δ*x*, Δ*y*, Δ*z* values revealed a larger motion of the M/E domain in mGBP2_holo_, can be explained with the fact that in mGBP2_holo_ this motion fluctuates around the initial structure involving considerable motions in opposite directions, whereas it evolves in only one direction in mGBP2_apo_. This results in smaller average *d*_480_ values in mGBP2_holo_. In both systems, the number of salt bridges takes on values between 0 and 3. More details are uncovered if one analyzes the distribution of descriptor combinations. For mGBP2_apo_, an interesting finding is that larger *d*_480_ values are preferentially found together with a larger RMSD_GC_, and that these motions are not correlated with the existence of the three salt bridges. In mGBP2_holo_, on the other hand, the RMSD_GC_ is not related to the motions of the M/E domain, which can be explained with the generally low mobility of the guanine cap following GTP binding. Furthermore, there is a fourth macrostate that was not sampled by mGBP2_apo_ and is characterized by the permanent formation of all three salt bridges, which results from the backbone shift of *α*4’ away from the nucleotide binding pocket following GTP binding, bringing it closer to the E domain (Fig.**3b/c**). While we identified states where none of the three salt bridges was present in either of mGBP2_apo_ and mGBP2_holo_, we did not sample the complete disruption of the G–E interdomain contact. One might expect this to see for mGBP2_holo_ as the complete disruption of the interdomain contact was observed for hGBP1 during the GTP hydrolysis reaction and shown to be needed for the formation of oligomers larger than dimers^44^. However, we neither simulated the hydrolysis, which might provide the energy needed to cause further motions, nor the transient state of the hydrolysis reaction, which may explain why we did not monitor the detachment of the E domain from the G domain. It might also occur on time scales beyond those tested in our simulations, despite applying an enhanced sampling method.

In summary, the TNs revealed further information about correlations between motions in mGBP2, yet allosteric communication pathways could not be uncovered yet. For this, non-equilibrium MD simulations would be better suited^45^, while enhanced sampling techniques other than HREMD, such umbrella sampling MD could be used to simulate the detachment of the E domain from the G domain. Such simulations will be pursued in our future studies of GBPs.

#### 2.1.5 Convergence tests

In order to test if the HREMD simulations involving 16 × 200 ns had indeed converged, we performed for mGBP2_holo_ another HREMD simulation employing 40 replicas of 400 ns each, amounting to 16 *μ*s in total for that simulation. As before, the analysis was performed for the target replica. We further checked how much conformational sampling was achieved by the standard 1 *μ*s MD simulation of the mGBP2_holo_ monomer. This question is of relevance for our simulations of the mGBP2 dimer and membrane-bound mGBP2, where the systems were too large for running HREMD simulations.

For both simulations we performed the same kind of analysis as done for the 16 × 200 ns HREMD simulation. In Supplementary Fig. S4**a** the RMSF values are shown. In both the 40 × 400 ns HREMD and the 1 *μ*s MD simulation the overall flexibility of the protein is very similar as in the initial HREMD simulation. The conformational space sampled by both HREMD simulations is in particular very similar, as confirmed by the RMSF values projected onto the protein (Supplementary Fig. S1**c** vs. S1**d**) and a similar number of clusters identified for the different motifs and loops of the G domain and the geranylgeranyl group (Supplementary Table S3). In the 1 *μ*s MD simulation, mGBP2_holo_ shows lower flexibility (Supplementary Fig. S1**e**) and also has slightly lower numbers of clusters (Supplementary Table S3). With regard to the hinge motion, this is stronger during the 40 × 400 ns HREMD simulation with 69.6 Å maximal displacement, whereas in the 1 *μ*s MD simulation that motion is somewhat reduced to a 44.6 Å amplitude (Supplementary Fig. S5**a/b**). We also performed a PCA and extracted the conformations corresponding to the minimal and maximal PC1 and PC2 values (Supplementary Fig. S5**c** and **d**). Finally, we monitored the motions of the M/E domain tip by measuring the Cartesian displacement coordinates of L480 with respect to its position in the mGBP2 homology model (Supplementary Fig. S5**e** and Table S2). In the 40 × 400 ns HREMD simulation, the M/E domain motions were of similar width as in the 16 × 200 ns HREMD simulation, yet shifted toward to more negative values in the x and the y direction. This is also confirmed by the PCA values with PC1_min_ of −14.7 nm^2^, but only PC1_max_ of 9.7 nm^2^, indicating that the hinge motion is more directed toward the *α*13 helix than in the shorter HREMD simulation of mGBP2_holo_. In the 1 *μ*s MD simulation, on the other hand, the hinge motion is less pronounced, as supported by the reduction of the sampled PC1 and PC2 space as well as the smaller Δ*x*, Δ*y*, and Δ*z* values (in absolute terms) for the movement of the M/E domain tip.

In conclusion, the computational cost of a 40 × 400 ns HREMD simulation is very high, while the afforded additional sampling of is minor. Thus, we concluded that the HREMD simulation with 16 replicates of 200 ns simulation was sufficient. Furthermore, the conformational space is better sampled by HREMD simulations, but the 1 *μ*s MD simulation shows similar results for the mGBP2_holo_. Therefore, we can use this time length for the MD simulations of the bigger systems.

### 2.2 Dimeric mGBP2 in solution

It was demonstrated that mGBPs accumulate as hetero- and homo-multimers, in vesicle-like structures in the cytosol and at the PVM, during host responses^7^. We therefore wanted to probe the dynamics of the mGBP2 dimer as the smallest multimer. We generated the mGBP2 dimer based on the crystal structure of the G domain dimer that was resolved for hGBP1 (PDB ID: 2BC9)^31^. The resulting dimer structure with monomer 1 and 2 (M1 and M2) is depicted in Fig. 1**b**. The use of AlphaFold-Multimer^46^ resulted in very similar mGBP2 dimer models that share the same interaction interface (Supplementary Fig. S6). We then simulated the dimer for 1 *μ*s. As mGBP2 dimerization was shown to preferentially occur in the presence of GTP, we modeled the protein in its GTP-bound state. We further included the geranylgeranyl group in the setup, considering that the dimer is expected to bind to the PVM via this lipid anchor. We thus modeled the protein as mGBP2_holo_ in our simulations of the dimer.

#### 2.2.1 Motions in the G domains and interactions in the dimer interface

We first compare the overall flexibility, as quantified by the RMSF values, of the mGBP2_holo_ in the dimer with that of the monomeric mGBP2_holo_, using the 1 *μ*s MD simulation of the monomer for the sake of comparability (Supplementary Fig. S4**b**). In general, the distribution of flexible and rigid protein regions is very similar in the mGBP2_holo_ monomer and dimer, which is also visible from the color-coded projections of the RMSF values onto the protein structures (Supplementary Fig. S1**e/f**). Only subtle differences are observed. As in the monomer, the two G domains of the dimer are stable, except for L1 and *α*4’ in M1. For the M/E domain, M1 only shows high flexibility at the tip (residues 415–430 and 468–504), whereas M2 exhibits a wholly flexible *α*12. To characterize the different motions, we executed a clustering with a cutoff value of 2.5 Å for the G domain’s structural elements (Fig.**6a/b**) and for the geranylgeranyl anchor with a cutoff value of 2.0 Å (Fig. 6**c**). The numbers for the resulting clusters and their populations are summarized in Supplementary Table S3. The clustering confirms that L1 (34 clusters) in M1 is more flexible than it is in M2 (15 clusters). This can be explained by the different influences of the geranylgeranyl anchor in M1 and M2. In M1, it interacts more with the L1 loop similar to the behavior in the other simulations of mGBP2_holo_ in solution, while in M2 the anchor causes an unfolding in the *α*12 (residues 523–544), creating a cavity where it does not interact with L1.

**Figure 6.**
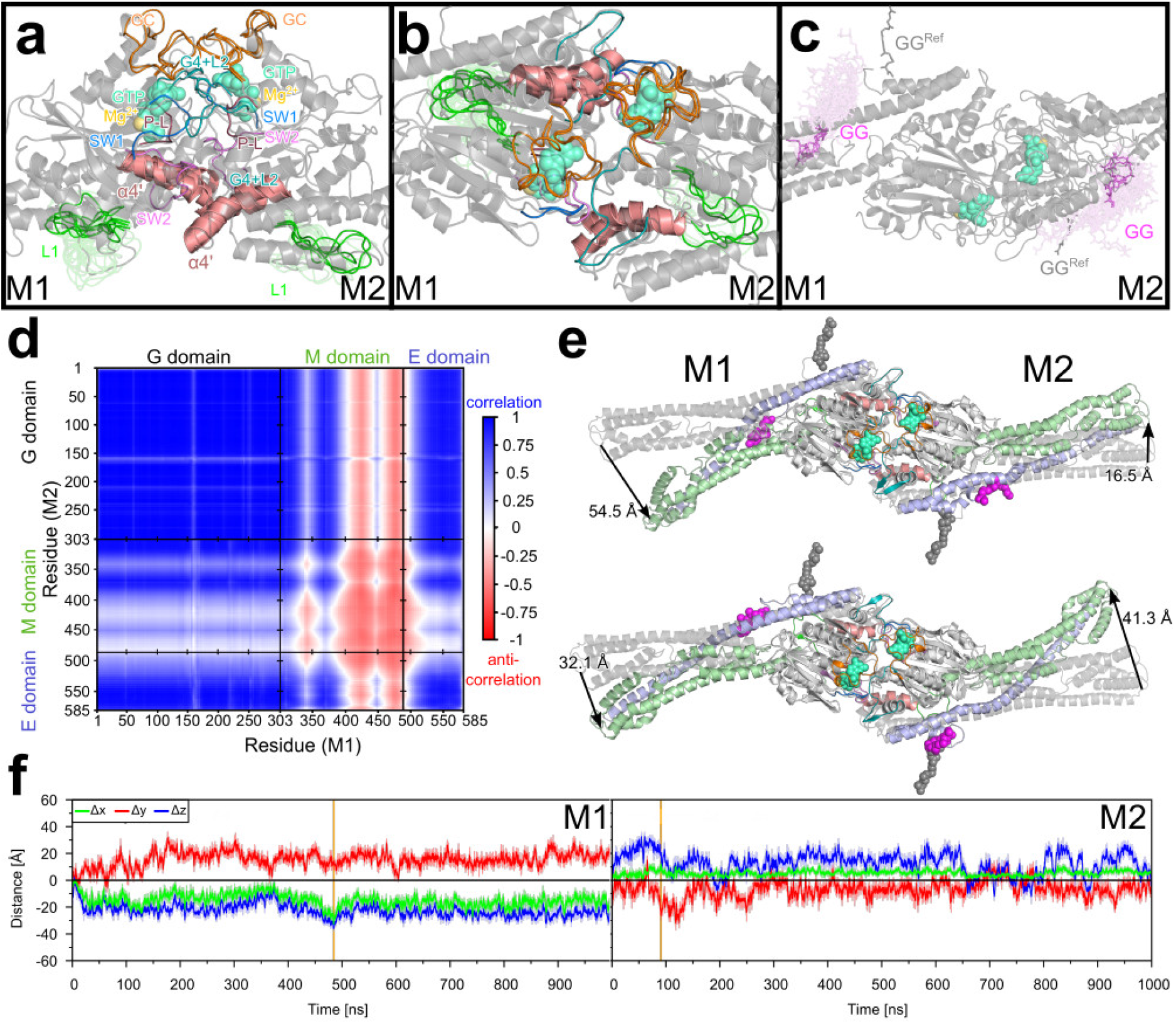
Molecular motions in the mGBP2_holo_ dimer obtained from a 1 *μ*s MD simulation. Conformational clusters of the G-domain motifs, the L1, and the guanine cap in the side view **(a)** and front view **(b)**. In **(c)** the clusters of the geranylgeranyl lipid anchor (GG) are shown. The same coloring scheme as in Fig. 3 was used. **(d)** The dynamic cross-correlation between the motions of the C_α_ atoms of M1 and M2 is shown. Blue and red colors represent correlation and anti-correlation, respectively. Uncorrelated motions, corresponding to the value 0, are shown in white, and correlation values of < |0.5| can be considered insignificant. The different domains are separated by black lines and the domain names given on the axes. **(e)** The largest deviations from the start structure as sampled in the MD simulation are depicted, and the amplitudes (in Å) for the tip of the M/E domain are given. **(f)** The motions of residue L480 relative to the start structure of the MD simulation are quantified in terms of Δ*x* (green), Δ*y* (red), and Δ*z* (blue) in M1 (left panel) and M2 (right panel) after aligning the trajectory on the G domain of protein M1 and using the same coordinate definition as shown in Fig. 5. The solid lines display the running average and the shades illustrate the raw data. The zero line is shown in black, whereas the vertical orange line indicates the frame with the maximal distance of the L480 residue between reference structure and the simulation.

Of special interest in the dimer is the guanine cap as it provides the interface between M1 and M2. In the dimer, both guanine caps have the same ordered structure as in the simulations of the mGBP2_GTP_ and mGBP2_holo_ monomer (Fig.**6a/b**). Also the numbers of clusters are similar, with two/three cluster conformations per guanine cap found in the dimer and two clusters identified in the 1 *μ*s simulation of the mGBP2_holo_ monomer (Supplementary Table S3).

To identify the key residues in the dimer interface, we calculated the contact probabilities with a cutoff value of 5 Å between the residues of M1 and M2 (Supplementary Fig. S7). The first observation is that several of the interactions between the two proteins are symmetric, such as Y47–Y199, E105–N190/H202 and D237–K243, which results from the symmetric structure of the homodimer. All special interactions like hydrogen bonds and salt bridges are summarized in the Supplementary Table S4, while the residues which are close to each other, yet without certain interactions are not listed. Some very stable hydrogen bonds are formed, which can be between backbones, side chains or between a backbone and a side chain. The two guanine caps are held together by stable salt bridges, like M1:D237–M2:K243, M1:E259–M2:R242/K243, and M1:K243–M2:E259 as well as a stable symmetric hydrogen bond between the side chain of R238 and the backbone of D182 in the G4+L2 motif. In the homologous hGBP1, the two arginine residues R240 and R242 are known to influence dimerization, as the protein remains a monomer if these two residues are mutated to alanine^37^. In mGBP2, the corresponding residues could be R238 and R242, but also K243, which form stable interprotein contacts. Furthermore, residues from the other G-domain motifs, in particular the phosphate-binding loop, SW1 and SW2, are also involved in the dimeric interface, where E105 (SW2) builds a stable hydrogen bond network with N190 and H202 (G4+L2) from the other monomer. The *α*4’ is also affected by dimerization, as it interacts with the SW2 motif of the other monomer via a salt bridge (K213–D108), a hydrogen bond (R221–V104), and a hydrophobic contact (F217–V104).

Taken together, numerous interactions are possible within the dimer interface of the mGBP2 homodimer. However, since for hGBP1 it was demonstrated that only two arginine residues in the guanine cap have a major impact on dimerization^37^, it can be speculated that the corresponding arginine or lysine residues R238, R242 and K243 play a similarly dominant role in mGBP2 dimerization.

#### 2.2.2 Large-scale motions of the dimeric mGBP2

To analyze the dynamics of the mGBP2 dimer, and whether the motions in the two proteins (M1 and M2) composing the dimer are correlated to each other, we determined the dynamic cross-correlation (DCC) for the motions of the C_*α*_ atom. The results in Fig.**6d** show that correlated and anti-correlated motions occur in a rather symmetric manner between the domains of the two proteins, which derives from the symmetry of the homodimer. The two G domains, the two E domains, as well as the G domain of one protein and the E domain of the other protein move in a correlated fashion. An exception is the loop L1 of both proteins (residues 151–160) whose motions are not, or only very little, correlated with the dynamics in the respective other protein. The motions of the two M domains are inversely correlated to each other. The relationship between the motions of the M domain with those of the G and E domains is mixed. The first half of the M domain up to about residue 400, which includes *α*7 (residues 310–340), *α*8 (348–369), and the first half of *α*9 (377–422), moves in a correlated manner with the G domain and E domain of the other protein in the dimer. The motions of the second part of the M domain, involving the second half of *α*9, *α*10 (430–447), and *α*11 (455–476), are inversely correlated with the motions of the G domain and E domain of the respective other protein. These anti-correlations are only weak to nonexistent for the motions between the M domain of M2 and those of the G and E domain of M1, whereas they are clearly present between the M domain of M1 and the G and E domain of M2. To understand these (anti-)correlations in the dimer dynamics, we elucidated what kind of motions take place. To this end, we monitored the motions of the tips of both M/E domains as they are the most flexible regions in the dimer according to the RMSF plot in Fig. S3b. In Fig. 6**e**, the snapshots corresponding to the largest deviations of the M/E domain relative to its starting structure, i.e., the homology model of the dimer, are shown, revealing a hinge motion similar to that taking place in the mGBP2 monomer. Protein M1 is seen to exhibit somewhat larger hinge motions than M2, with a maximal deviation of residue L480 from it starting structure of 54.5 Å in M1, as compared to 41.3 Å in M2. Fig.**6e** further suggests that the hinge motions in M1 and M2 are anti-correlated with respect to each other, as a large (small) amplitude in M1 is recorded together with a small (large) amplitude in M2. Next, we computed the Cartesian displacements of the L480 residues to quantify the hinge motion, which can be seen in Fig.**6f** and are summarized in Supplementary Table S2. As indicated by the snapshots in Fig.**6e** and by the DCC analysis in Fig. 6**d**, the hinge motions in M1 and M2 are anti-correlated to each other. In particular, the hinge motion of M1 takes place in positive *y*-direction, whereas M1 moves mainly into the opposite *y*-direction. Interestingly, in the dimer the two M/E domains also exhibit considerable motions along the *z*-axis, again into opposite directions (negative *z*-direction in M1, positive *z*-direction in M2), which did not occur in the monomer (Fig. 5**b**). The same applies to motions along the *x*-axis, which are however less pronounced than those along *y* and *z*. In general, the amplitudes of the motions occurring in M2 are smaller than those in M1, which agrees to the findings from the snapshots in Fig. 6**e**.

To summarize our observations of the dimeric state, the two guanine caps are rather rigid and stabilizing each other. This in turn may help to hold GTP more strongly in its binding site and is thus more efficiently hydrolyzed, as observed in experiments^7^. The hinge motion of the M/E domains is also affected by dimerization, as it moves in somewhat other directions than in the monomer. Moreover, the two hinge motions occurring in M1 and M2 of the dimer are anti-correlated. We elucidated the most relevant interprotein interactions that stabilize the dimer. Via mutations studies, it could be checked if any of these interactions, in particular those involving R238, R242 and K243, are indeed necessary for the dimerization process.

### 2.3 Membrane binding of mGBP2

Next, we analyzed the binding of mGBP2 to lipid membranes. Since the lipid composition of the PVM of *T. gondii* is not known yet, we first performed experiments to identify the lipids to which mGBP2 prefers to bind, and then used this information to repeat the simulation of membrane-bound mGBP2 using the same lipid composition as studied experimentally and to which mGBP2 bound well, which is 75% 1,2-dioleoyl-sn-glycero-3-phosphocholine (DOPC), 20% cholesterol, and 5% cardiolipin. With regard to the simulations of membrane-bound mGBP2, we started by analyzing the behavior of mGBP2 on a pure POPC (1-palmitoyl2-oleoyl-sn-glycero-3-phosphocholine) bilayer serving as a reference system, and then continued with a biologically more relevant mixed lipid bilayer using the lipid composition determined experimentally.

#### 2.3.1 Dynamics of mGBP2 on a POPC membrane

We first conducted 1 *μ*s MD simulations of mGBP2 on a POPC membrane starting from the structure shown in Fig.**1c** and considering mGBP2 without and with GTP being present. For the membrane binding, the geranylgeranyl lipid anchor was always attached to the protein and inserted into the membrane. In order to account for the presence of the geranylgeranyl anchor, but absence of GTP, we denote this system as 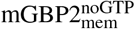, while 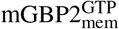 is used for mGBP2 featuring both bound GTP and the geranylgeranyl anchor. We are aware that the 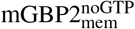 system does not reflect reality as GTP is needed for mGBP2 to bind to a membrane^10^. However, by comparing 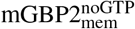 with 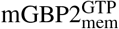 we are able to investigate the influence of GTP on the interplay between mGBP2 and the membrane.

As for the protein in solution, we find that the binding of GTP to membrane-bound mGBP2 decreases the flexibility of the G domain motifs G4+L2, SW1 and the guanine cap. This can be seen from the RMSF values in Supplementary Fig. S4**c** and from the reductions in the numbers of clusters when applying conformational clustering to the various motifs and loops (Supplementary Table S3). This applies in particular to the SW1, G4+L2 motif, and the guanine cap, which are very flexible in the 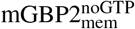 system, with 34, 62, and 101 clusters, respectively. These numbers reduce to 4, 11, and 5 clusters upon GTP binding in the 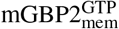, which are similar, albeit somewhat higher, as in the monomeric mGBP2_holo_ in solution. With regard to the movements of the M/E domain, we find that it is still flexible, despite the direct neighborhood of the lipid membrane, as can be seen from the RMSF values (Supplementary Figs. S1**g/h** and S5). The hinge motion can still take place on the membrane (Fig. 7a) and is not much affected by it, as assessed by the motions of the tip of the M/E domain. Using L480 as measuring point, as before, the maximal deviation from its starting structure is 57 Å for 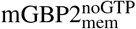 and 59 Å for 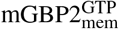, and is around 25 Å on average in both systems. In the 1 *μ*s MD simulation of mGBP2_holo_ in solution, very similar values were reached.

**Figure 7.**
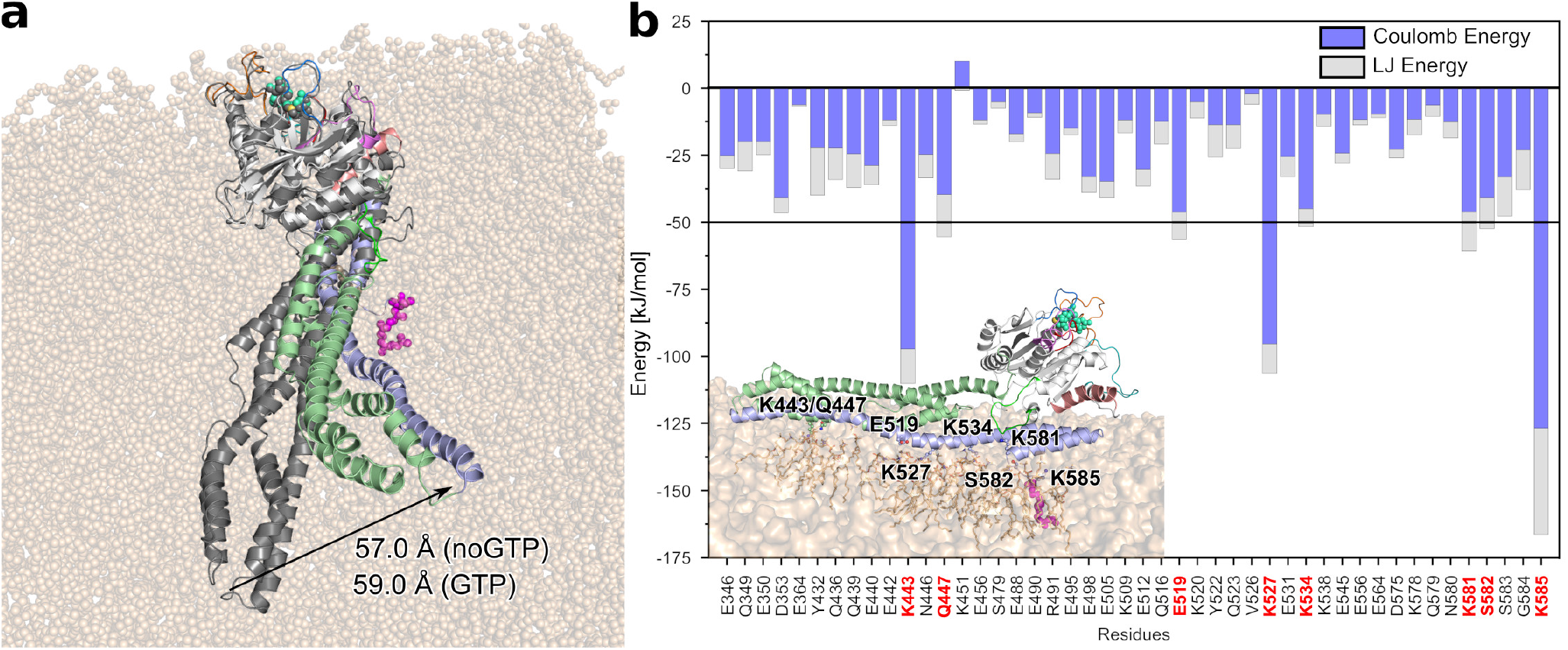
Hinge motion and energies of the strongest interactions between mGBP2 residues and the POPC membrane. **(a)** The conformation with the highest distance to the start structure, as measured by the distance of L480 with respect to the MD start structure (in gray), is shown and the distances for 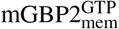 and 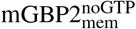 are given. During the motion the tip of the M/E domain detaches from the membrane surface. **(b)** The 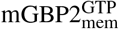–membrane energies are decomposed into Coulomb (blue) and Lennard-Jones (gray) contributions. The residues with a total interaction below –50 kJ/mol (horizontal black line) are highlighted with red labels and considered as key residues for interacting with the membrane. They are all part of the M/E domain, as can be seen in the structure figure in the lower left corner.

To characterize the motions of the protein relative to the membrane, we calculated the change in the *z*-position, which corresponds to the bilayer normal, between the center of mass of selected parts of the protein and that of the phosphor atoms of the lipids of the upper bilayer leaflet. Supplementary Fig. S8**a** shows the definition of the coordinate system as well as the parts of the protein used for the Δ*z* calculation. These include three residue areas from the G domain, namely L1 (151–161), G4+L2 (181–196), and the guanine cap (235–256), the whole M domain (303–476) and only the M/E tip (480), as well as *α*12 and *α*13 of the E domain, and finally the geranylgeranyl lipid anchor. The Δ*z* values of these regions are plotted in Supplementary Fig. S8**b** and **c** for 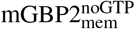 and 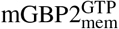, respectively. A statistical analysis is provided in Supplementary Table S5. The first observation is that 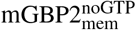 and 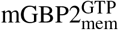 exhibit no big differences in their motions. Secondly, no dissociation of the E domain from the membrane is observed. Thirdly, with regard to the G domain one finds that in both systems, the loop L1 is generally closer to the membrane than G4+L2 and the guanine cap are. However, even L1 does not interact with the membrane, as its distance from the membrane surface is almost always 20 Å or higher. The center of mass of the M domain has also no membrane interactions, while the tip of the M/E domain touches recurrently the membrane. The repeated contact formation with the membrane surface and loss thereof are a result of the hinge motion of the M/E domain. However, the contacts formed between the membrane and both helices of the E domain are less affected by the hinge motion, as they are close to the membrane most of the time (fluctuating around 10–11 Å, Supplementary Table S5). Especially *α*13 interacts intimately with the membrane. The geranylgeranyl lipid anchor lies under the zero line, meaning that it remains fully membrane-inserted throughput the whole simulation. The similarity of the motions of 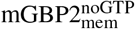 and 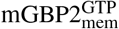 relative to the membrane suggests that GTP has no direct effect on the mGBP2-membrane interactions.

In order to elucidate the residues that contribute the most to the interaction with the POPC membrane, we calculated for the 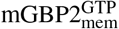 system the protein-membrane interaction energy and decomposed it into Lennard-Jones and Coulomb energies, *E*_LJ_ and *E*_Coul_. In Fig. 7**b**, these energies are plotted for all residues that feature a considerable interaction with the lipids. These residues belong all to the M/E domain, while the whole G domain shows no interaction with the membrane, although some of the areas come close. In particular, the lysines K443, K527 and K585 show the highest energies (with values below –90 kJ/mol; negative energies indicating attraction), together with Q447, E519, K581 and S582 (below –50 kJ/mol). These residues, especially the positively charged ones, interact with the head group of the POPC lipids, in particular with their negatively charged 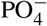 groups, giving rise to substantial electrostatic energies. The LJ interactions, on the other hand, are generally weaker and only appear together with electrostatic interactions, and not on their own. This indicates that, apart from the geranylgeranyl lipid anchor, none of the other parts of mGBP2 inserted into the membrane but remained on its surface. It was already demonstrated for mGBP2 and hGBP1 that the lipid anchor is necessary for the accumulation of the proteins at the PVM or GUVs^7,8^. From the current analysis we conclude that in addition to the lipid anchor, also some of the lysine residues of the M/E domain are strongly involved in the mGBP2-membrane interactions.

In summary, mGBP2 is stable on the POPC membrane, with little to no effects caused by the presence of GTP. The protein can still perform its characteristic hinge motion. However, this motion has no noteworthy effect on the membrane properties.

#### 2.3.2 Identification of mGBP2’s preferences for lipid binding

Considering that a pure POPC membrane is of limited biological relevance, the aim was to repeat the simulations with a membrane containing different components. However, the compositions of the T. gondii plasma membrane and the PVM are generally ill-defined. To gain first insights into potential mGBP2–lipid interactions, a protein–lipid overlay assay was used. Membrane lipid strips are hydrophobic membranes containing 100 pmol spots of fifteen different membrane lipids, which we incubated with 1.25 *μ*g GFP-mGBP2 protein or single GFP as control. The quantification of the protein–lipid overlay assay revealed a strong and significant binding of GFP-mGBP2 to cardiolipin (CL), among other lipids that will be analyzed elsewhere. Cardiolipin represents a mitochondria-specific phospholipid, where it is mainly located in the inner membrane (up to 20% of total lipids)^47^. The control, GFP alone, did not bind to any membrane lipid spots on the lipid strips (data not shown). To further corroborate these findings, confocal microscopy applied to GUVs was used to observe the direct interaction of GFP-mGBP2 with membranes in real time. The GUVs were prepared from a membrane basic structure composed of DOPC, cholesterol, CL, and TexasRed. As a control, GUVs containing phosphatidylserine (PS) instead of CL were used, since no binding of GFP-mGBP2 to PS could be observed on membrane lipid strips. Co-incubation of GFP-mGBP2 with the CL-GUVs showed a clear accumulation of mGBP2 at the synthetic membranes, while no binding of mGBP2 to PS-containing GUVs was visible (Fig. 8**a**). Thus, using two different techniques, we identified CL as a phospholipid being specifically targeted by GFP-mGBP2. Moreover, the comparison between the CL and PS results indicates that a negatively charged lipid headgroup is not sufficient to induce membrane binding of mGBP2.

**Figure 8.**
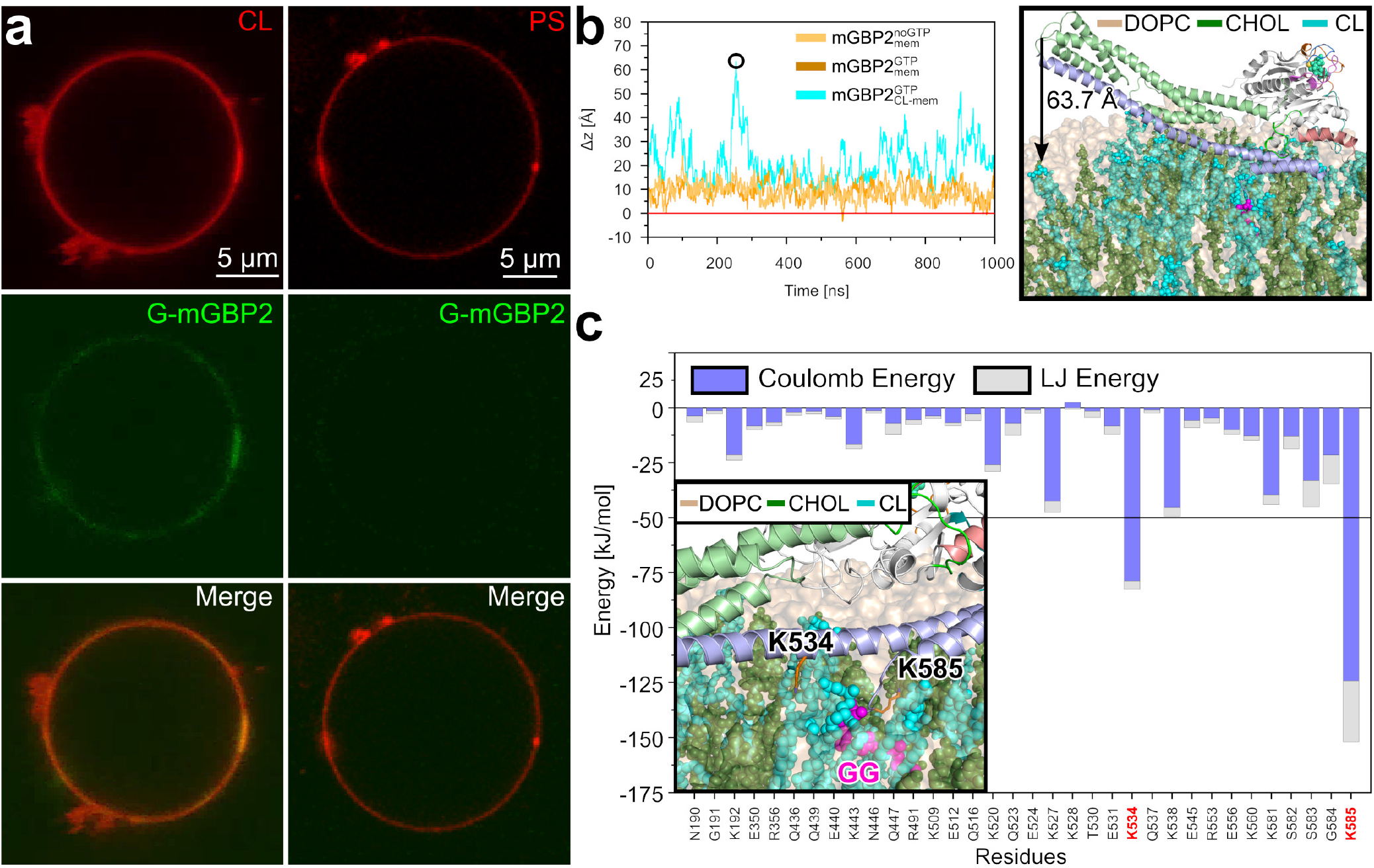
The binding of mGBP2 to mixed lipid membranes. **(a)** GFP-tagged mGBP2 accumulates at TexasRed-labeled GUVs containing cardiolipin (CL, left) but not at GUVs with phosphatidylserine (PS, right). **(b)** Simulations of mGBP2 bound to a DOPC/cholesterol/cardiolipin bilayer revealed large-scale motions of the M/E domain, as monitored by the distance (Δ*z*) between residue L480 and the average position of the lipid head groups (red line). The results for the 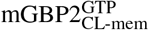 system (cyan) are compared with the motions found for 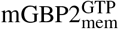 (brown) and 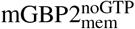 (orange). The snapshot on the right shows the conformation with the largest movement of the M/E domain (Δ*z* = 63.7 Å), where the lipids are colored in ochre for DOPC, green for cholesterol, and cyan for cardiolipin. **(c)** The time-averaged residue–membrane interaction energies are decomposed into *E*_Coul_ (blue) and *E*_LJ_ (gray). The residues with a total interaction below –50 kJ/mol (black line) are highlighted with red labels and considered as key residues for interaction with the membrane. They are both part of the E domain, as can be seen in the structure figure on the left.

#### 2.3.3 Dynamics of mGBP2 on a cardiolipin-containing membrane

Using the experimental results as input, we performed a 1 *μ*s MD simulation of GTP-bound mGBP2 pre-inserted into a membrane containing DOPC, cholesterol, and cardiolipin in a ratio of 75%/20%/5%, mimicking the lipid composition of the GUV study. This system is called 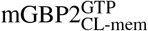 in the following. As for the simulation with the pure POPC membrane, we measured the motions of residue L480 relative to the membrane surface, denoted as Δ*z* in Fig. 8**b**. It can be seen that mGBP2 is much more mobile on the mixed membrane than on the pure POPC membrane; on the CL-containing membrane the tip of the M/E domain can move up to 64 Å away from the membrane surface (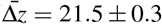 Å mean ± standard error of the mean), while the geranylgeranyl group at the C-terminal end of the E domain remained stably inserted in the membrane. The corresponding 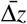 values for the pure POPC-bilayer systems are 9.3 ± 0.1 Å for 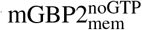 and 8.9 ± 0.1 Å for 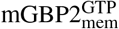. To reveal the source of the larger mGBP2 motions on the mixed membrane, we calculated the interaction energies, decomposed into *E*_Coul_ and *E*_LJ_, between mGBP2 and the different lipids (Supplementary Fig. S8**d**) and also split the energies into per-residue contributions (Fig. 8**c**). Compared to the 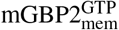 system, in binding of mGBP2 to the mixed membrane the negatively charged residues E350, E440, E512, and E531 and the polar residues Q436, Q437, Q477, S582, and S583 are less involved, which can be explained by the presence of the negatively charged CL lipids. Two lysine residues of 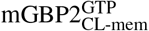, on the other hand, play important roles in mGBP2–membrane binding, reaching interactions energies of below –50 kJ/mol: K534 and K585. They interact with the headgroup of CL, as the inset of Fig.**8c** and Supplementary Fig. S8**e** (snapshot at 551 ns) show. The latter figure also reveals that the geranylgeranyl anchor forms contacts with the fatty acids of CL and the ring system of cholesterol (snapshots at 868 and 962 ns). Representative interactions involving the G and M domain are shown for N190 and K192 of the G4+L2 motif being in contact with the hydroxy group of cholesterol (snapshot at 88 ns), and K443 of the M domain interacting with CL (snapshot at 868 ns). Next, we study the evolution of the interaction energies of mGBP2 with the different lipids (Supplementary Fig. 8**d**) and compare it with the motion of L480 relative to the membrane surface (Fig. 8**b**). As expected, the interaction energies are highly negative when the tip of the M/E domain is close to the membrane, indicating tight binding of mGBP2 to the membrane. Surprisingly, the mGBP2 motion away of the membrane is not caused by strong, but only small repulsions between the protein and the membrane surface, in particular with DOPC. For instance, large Δ*z* values between 200 and 300 ns are preceded by electrostatic repulsion between mGBP2 and DOPC, which is not completely negated by attractions between CL and the protein. However, not all motions away from the membrane can be time-correlated with these interactions, yet the negatively charged residues in the E domain close to L480 play a role in causing the Δ*z* motions (Supplementary Fig. S8**d**).

To sum up, the simulations of mGBP2 being bound to a DOPC/cholesterol/cardiolipin bilayer revealed that the negatively charged cardiolipin induced considerably larger motions of the M/E domain on the membrane than on the zwitterionic POPC membrane. Nonetheless, also in that case no changes in membrane curvature were recorded. It will be interesting to study what the effects of mGBP2 oligomers on a membrane are.

## 3 Discussion and conclusion

In this study, we provide insights into the conformational changes of mouse GBP2, the orthologue of human GBP1. We modeled mGBP2 as a monomer in GTP-free and GTP-bounds states, as a dimer, as well as anchored to different lipid membranes, using atomistic molecular dynamics simulations and an enhanced sampling method, Hamiltonian replica exchange molecular dynamics. This is the first report of a GBP being simulated in the GTP-bound state, in addition to considering its isoprenylation. For setting up the mGBP2–membrane systems, we identified the lipids that mGBP2 prefers for binding using membrane lipid strips and GUV experiments, which were then included in our simulations.

We find that GTP binding considerably stabilizes the G domain of mGBP2. Even the guanine cap, which is highly flexible without GTP, becomes more ordered and less flexible upon GTP binding. In the biochemical study of Kravets et al.^10^, the GTP binding site was characterized in detail with the K51A and R48A mutants affecting phosphate binding, and E99A in the SW2 and D182A in the G4 motif influencing GTP activity and oligomerization. Compared to hGBP1 with its previously reported dissociation constant *K*_D_ of 1.1 *μ*M^48^, mGBP2 has a 2.4-fold higher affinity to GTP with a *K*_D_ of 0.45 μM^10^. This difference might be explained by their slightly different amino acid sequences. Upon GTP binding, the key residues of the mGBP2’s G domain interact with the nucleotide, as was previously demonstrated for hGBP1^32,48^. The decreased flexibility of the guanine cap following GTP binding seen here for the mGBP2 can explain why this shifts the monomer–dimer equilibrium to the dimeric state, like in hGBP1^28,37,49^, as this should reduce the penalty for dimer formation arising from the loss of conformational entropy of the guanine cap. In the dimeric interaction interface, we identified two residues in the guanine cap, R238/R244, which could have the same function as R240/R244 in the human orthologue^37^.

The characteristic hinge motion that we previously identified in hGBP1^35^ was seen in all states of mGBP2 that we simulated. Interestingly, this hinge motion may lead to a closed state. Having both a closed and open state is a property that was identified for other dynamin-like proteins, including the human myxovirus resistance protein and also the bacterial dynamin-like protein^50–52^. In solution, the hinge motion of mGBP2 can cover up to 70 Å whereas this decreases to only 30 Å in the dimeric state or on the membrane. As we know, that different mGBPs localize at the PVM of *T. gondii* as either dimer or higher multimer states, the hypothesis arises that this hinge motion can cause membrane damage, whereupon the membrane might be breaking with sufficient amount of GBPs. This has been shown to be the case for bin-amphiphysin-rvs (BAR) domain proteins^53,54^. However, with only one mGBP2 molecule being bound to the membrane, we did not observe any significant changes in the membrane properties, such as no strong curvature, regardless of whether GTP was present or not, and regardless of the composition of the membrane. We considered both a pure POPC membrane and a mixed membrane containing 75% DOPC, 25% cholesterol, and 5% cardiolipin. For GUVs of the latter composition we demonstrated that mGBP2 assembles at the membrane surface, and in preceding membrane lipid-strips experiments we had identified cardiolipin as one of the lipids with the highest binding potential for mGBP2. This finding extends the previously reported interactions of hGBPs with lipopolysaccharides (LPS) and provides more evidence that GBPs can directly interact with pathogen associated molecular patterns^55,56^. Interestingly, mGBP2 did not bind to membranes containing phosphatidylserine. This suggests that despite the several basic residues, especially lysine residues, that are found in helices *α*12/13, the charge attraction between these residues and the negatively charged lipids is not sufficient to explaing the lipid-binding specificity of mGBP2. The simulations revealed different protein dynamics on the zwitterionic POPC and anionic cardiolipin-containing membrane. On the latter, mGBP2 was considerably more dynamic, with a large-scale up and down motion of the M/E domain on the membrane, while the C-terminal of the E domain and the geranylgeranyl anchor remained membrane-bound. This motion is caused by the general flexibility of the M/E domain, showing itself as permanently present hinge motion, and electrostatic repulsion between lysine residues close to the tip of the M/E domain and the anionic surface charge of the membrane. Nonetheless, this protein motion is not strong enough to cause changes in the membrane curvature or other membrane damage. We expect that membrane-bound mGBP2 multimers are needed to cause membrane damage^57,58^, which will be addressed in our future work.

In recent studies, it was demonstrated that hGBP1 can polymerize by detaching the E domain from the other two domains to swing it out, giving rise to disk-like structures, where the G domains are at the outside of the disk and the farnesyl groups assembled together inside the disk^8,59^. A conformation with the E domain in the folded-out position and dimerization via the G domains, as described here, could also tether two giant unilamellar vesicles together via the farnesyl lipid anchor^8,18,39^. Until now, a similar folding out of the E domain has not be shown yet for mGBPs. Nonetheless, such a motion should be considered in future simulations studies of mGBP2 too. It should be noted that the dimer state studied here should have a hard time to insert both geranylgeranyl lipid anchors into the membrane at the same time as they are oriented into different directions. Also this aspect needs attention in the future.

## 4 Methods and materials

### 4.1 Molecular dynamics simulations

For the MD and HREMD simulations, the Amber99SB*-ILDNP force field^60–62^, TIP3P water^63^, and GROMACS 2016^64,65^ were applied. Only for the simulation involving the mixed lipid membrane we used the CHARMM36 force field^66, 67^ as it provides force-field parameters for cardiolipin^68,69^. All simulations were run at a temperature of 310 K (37°C) and a pressure of 1 bar. In Table 2, all simulations performed in this study are summarized, amounting to a total simulation time of 31.6 *μ*s. Detailed model preparation and simulation descriptions can be found in the **Supplementary Information**.

**Table 2.** Summary of simulations performed in this work.

| Simulation | System | Size in atoms | Replicas/Runs × Length | Cumulated time |
| --- | --- | --- | --- | --- |
| HREMD | mGBP2<sub>apo</sub> | 131,493 | 16 × 200 ns | 3.2 μs |
| HREMD | mGBP2<sub>GTP</sub> | 129,116 | 16 × 200 ns | 3.2 μs |
| HREMD | mGBP2<sub>holo</sub> | 130,665 | 16 × 200 ns | 3.2 μs |
| HREMD | mGBP2<sub>holo</sub> | 130,665 | 40 × 400 ns | 16 μs |
| MD | mGBP2<sub>apo</sub> | 129,246 | 1 × 1 μs | 1 μs |
| MD | mGBP2<sub>holo</sub> | 130,665 | 1 × 1 μs | 1 μs |
| MD | mGBP2<sub>holo</sub> dimer | 573,673 | 1 × 1 μs | 1 μs |
| MD | mGBP2<sub>mem</sub><sup>noGTP</sup> | 659,946 | 1 × 1 μs | 1 μs |
| MD | mGBP2<sub>mem</sub><sup>GTP</sup> | 662,603 | 1 × 1 μs | 1 μs |
| MD | mGBP2<sub>CL-mem</sub><sup>GTP</sup> | 664,034 | 1 × 1 μs | 1 μs |
| **Total simulation time** |  |  |  | **31.6 μs** |

### 4.2 Analysis of the simulations

If not stated otherwise, all analyses were performed using GROMACS 2016. For the analysis of the HREMD simulations, only the data collected by the target replica was used.

#### Flexibility

To describe the stability and flexibility of mGBP2, the RMSF of the C_α_ atoms around their average positions was determined for each residue. The RMSF describes the positional change of the selected atoms as time-average. A residue with an RMSF value over 2 Å was considered as flexible. We also projected the RMSF values onto the start structure using a color code, where red colors indicate highly flexible regions (≥ 2 Å) and blue colors are for rigid residues.

#### Secondary structure analysis

To calculate the average helix content of the E domain (*α*_E_) averaged over the whole trajectory, the DSSP (Define Secondary Structure of Proteins) algorithm^70^ was employed. It determines the secondary structure for all residues, from which the *α*-helical content of the E domain was determined.

#### Clustering analysis

Conformational clustering was performed to identify the most populated conformations of the G domain motifs and loops and the geranylgeranyl group. To this end, the algorithm of Daura et al.^71^ was applied to the C_*α*_ atoms of the structural element in question. This algorithm is based on the root mean square deviation (RMSD) between all structures collected in a trajectory, and an RMSD cutoff value of 2.5 Å was used to define cluster membership for the G domain motifs and loops. Prior the calculation, the trajectories were fitted onto the G domain of the starting structure. A similar clustering was performed for the geranylgeranyl lipid anchor, with the differences that **(i)** the trajectory was fitted using the whole protein and **(ii)** a cutoff value of 2 Å was used.

#### Hinge motion analysis

The main structural changes and movements of the M/E domain were identified based on a principal component analysis (PCA). For being able to compare the motions of the different mGBP2 systems within the same principle component (PC) space, we applied the PCA to the concatenated target replicas of the HREMD simulations (16 × 200 ns) of mGBP2_apo_, mGBP2_GTP_, and mGBP2_holo_. This step involved the removal of not only the solvent and ions, but also of GTP and Mg^2+^ from both mGBP2_GTP_ and mGBP2_holo_ and in the case of the latter, also the geranylgeranyl lipid anchor was removed, as the same number of atoms is needed for a joint PCA. Moreover, the trajectories were aligned to the G domain of the MD starting structure. Of the resulting principal components, the two main motions described by PC1 and PC2 were analyzed in detail. This included to separately project the target replica trajectories of the three HREMD simulations onto the space defined by PC1 and PC2. A PCA was also done for the additional simulations performed for mGBP2_apo_, i.e., the target replica trajectory of the 40 × 400 ns HREMD simulation and the 1 *μ*s MD simulation.

#### Transition networks

The TNs were calculated using the ATRANET (Automated Transition Network) software package^40,42^, developed by our group, and visualized with Gephi 0.9.7^72^. We defined the states of the network by three descriptors: (i) The presence of the three saltbridges R225–E554, R225–E561, and K226–E573 between G and E domain; this descriptor can adopt discrete values from 0 to 3. A salt bridge (SB) was considered to be present if the distance between the two atoms involved is below a cutoff value of 5.5 Å. (ii) The RMSD of the guanine cap’s C_α_ atoms (RMSD_GC_) in discretized intervals of 0.5 Å. (iii) The distance of the C_α_ atom of L480 compared to the starting structure in discretized intervals of 1 Å (d_L480_). The states are writte as [SB, RMSD_GC_, *d*_L480_]. We calculated the local communities of highly interconnected states using the modularity class feature of Gephi, allowing us to identify macrostates for mGBP2_apo_ and mGBP2_GTP_.

#### Dynamic cross correlation

To characterize the influence of the two proteins composing the dimer on each other’s motions, we applied a dynamic cross-correlation (DCC) as available in MD-TASK^73^ (https://md-task.readthedocs.io/en/latest/corr.html). This allows to determine the dynamic correlation between the atoms of a system, i.e., to what degree the atoms move together and whether their motion is in the same of the opposite direction. This analysis generated an *N* × *N* matrix, where *N* is the number of the C_*α*_ atoms, and each element (*i, j*) corresponds to the dynamical cross correlation between atoms *i* and *j*. This matrix was plotted using a color code, with blue colors representing correlations with values between 0.1 and 1 (complete correlation) and red colors indicating the anti-correlation with valued from −0.1 to −1 (complete anti-correlation). No or only very weak correlation is given, if the value is 0 ± 0.1, as represented by white in the DCC plot, yet all correlation values < |0.5| can be considered as insignificant.

#### Protein-protein interactions in the mGBP2 dimer

For the characterization of the mGBP2 dimer interface, all interprotein residue pairs where the two residues were within 12 Å of each other at any time during the MD simulation were identified, resulting in a 149 × 149 matrix. To further separate the strong residue-residue contacts from the weaker ones, the time-averaged contact probability was calculated. To this end, for each of the 149 × 149 residue pairs it was for every 1 ns determined if the pair in question was indeed in contact, using a 5 Å distance cutoff for the minimum distance between the residues to define the presence of a contact.

#### Analysis of the mGBP2 monomer on the POPC membrane

We monitored the motions of the three mGBP2 domains with respect to the POPC membrane by calculating the distance of the centers of mass of the three domains with respect to the membrane surface along the z coordinate (representing the membrane normal). To define the membrane surface position, the average position of the phosphor atoms of the POPC lipid headgroups were used. To further specify which parts of the domains are closest to the membrane, we also calculated the distances of the G domain motifs L1 (151–161), G4+L2 (181–196), and guanine cap (235–256), of the tip of the M/E domain (L480), of *α*12, *α*13, and the geranylgeranyl lipid anchor from the membrane surface. This analysis was conducted for every 1 ns. To identify the residues with the strongest interactions with the membrane, we calculated the time-averaged residue-membrane interaction energies *E*_LJ_ for the Lennard-Jones interactions and *E*_Coul_ for the electrostatic interactions.

#### Figures

For the generation of 3D protein structure figures we used PyMol^74^.

### 4.3 Experimental methods

#### Cell culture

mGBP2-/- murine embryonic fibroblasts (MEFs) stably expressing N-terminally GFP-tagged murine GBP2 (GFP-mGBP2) or GFP alone^11^ were cultured in Dulbecco’s modified Eagle’s medium (DMEM, Invitrogen/Gibco) supplemented with 10% (v/v) heat-inactivated low endotoxin fetal bovine serum (FBS, Cambrex), 2 mM L-glutamine (Biochrom) and 0.05 mM *β*-mercaptoethanol (Invitrogen/Gibco).

#### GFP-Trap^®^

The one-step immunoprecipitation of GFP-tagged proteins is performed by anti-GFP VHH coupled agarose beads (90 μm) (Chromotek) with a binding capacity of 3-4 *μ*g/10 *μ*l GFP-Trap® slurry. For one immunoprecipitation reaction, 10^6^ MEF cells expressing GFP-tagged mGBP2 were used.

#### Lipid-coated Strips

Membrane lipid strips (P-6002, Echelon Biosciences) were incubated with 1.25 μg isolated and purified GFP tagged mGBP2 proteins and single GFP-protein in 3% BSA for 2 h at room temperature. Lipid Strips analyses were performed with GFP specific first antibodies and detected with a goat anti-mouse HRP-linked secondary antibody (BD) and ECL reagent (GE Healthcare, Munich, Germany).

#### Giant unilammelar vesicle preparation

Giant unilammelar vesicles (GUVs) were prepared by electroformation (11Hz, 2V for 3 h at room temperature) using a construction scaffold of 74.75 mol % DOPC (1,2-Dioleoyl-sn-glycero-3-phosphocholine), 20 mol % cholesterol, 0.25 mol % TexasRed and 5 mol % lipid of interest (cardiolipin solved in chloroform). Lipid extracts were used from Avanti Polar Lipids.

#### Confocal microscopy

Confocal microscopy of interaction studies between isolated proteins and GUVs were performed using a LSM780 confocal microscope (Zeiss, Germany). GUVs solved in sucrose buffer were transferred in cellview cellculture slides (#543078, Greiner Bio-One, Germany) containing 250 *μ*l PBS. Image analysis was performed with the software ZEN (Zeiss).

## Acknowledgments (not compulsory)

We thank Dr. Bogdan Barz and Wibke Schumann for fruitful discussions.

## Author contributions statement

Conceptualization: B.S., D.D., K.P.; data curation: J.L., X.W., N.S.; formal analysis: J.L., N.S., M.S.; funding acquisition: B.S., D.D., K.P.; investigation: J.L., X.W, B.S., N.S., D.D., K.P.; methodology: J.L., B.S; project administration: B.S., D.D., K.P.; software: J.L., M.S.; supervision: B.S., D.D., K.P.; validation: J.L., B.S.; visualization: J.L., M.S., N.S.; writing – original draft: J.L., N.S., D.D.; writing – review & editing: B.S., D.D., K.P.

## Funding

Funding for this project was received from the Deutsche Forschungsgemeinschaft (German Research Foundation, https://www.dfg.de/) through Grant 267205415 (CRC 1208, Projects A06 and A07 (to D.D., K.P., and B.S.), PF 259/8-2 (to K.P) and the Manchot Foundation (to D.D. and K.P.). We gratefully acknowledge the computing time granted through JARA-HPC (project JICS6A) on the supercomputer JURECA at Forschungszentrum Jülich, through the Leibniz Supercomputing Centre (LRZ) of the Bavarian Academy of Sciences on the supercomputer SuperMUC-NG (project pn98zo), and through the Centre for Information and Media Technology at Heinrich Heine University Düsseldorf.

## Competing interests

The authors declare no competing interests.

## Additional information

### Supplementary information

Detailed description of the methods employed, supplementary figures S1 to S8, supplementary tables S1 to S5.

### Data availability

The datasets (trajectory files) created and analyzed during the current study are available from the corresponding author on reasonable request.

## Notes

### Competing Interest Statement

The authors have declared no competing interest.

### Summary of Updates

Experiments were done to further elucidate the mGBP2-membrane interactions. The results of these experiments resulted in novel simulations to be performed, involving a membrane composed of thee lipid types. To better unravel the domain-domain communication during the protein movements, transition networks were developed.

