## Supplementary Tables and Figures for "Domain motions, dimerization, and membrane interactions of the murine guanylate binding protein 2"

### Methods

#### Preparation of mGBP2 models

If not stated otherwise, the Amber99SB*-ILDN force field[1–3] was used for modeling mGBP2 and TIP3P for modeling water[4].

**GTP-free mGBP2 model.** The structure of mGBP2 was derived by Kravets et al.[5] applying homology modeling with hGBP1 in its nucleotide-free state (PDB ID: 1DG3)[6] as template using the SWISSMODEL server[7,8] (`https://swissmodel.expasy.org/`). The sequence identity between hGBP1 and mGBP2 is 68.4% as determined by T-coffee[9,10]. This resulted in a structural model of mGBP2 covering most of the amino acids 6 to 577 (out of 589 residues). The residues of the loop 1 (L1; residues 149–164) as well as some of the N- and C-terminal amino acids, including the CaaX sequence, are missing as they were not resolved in the hGBP1 structure used as template[5]. The missing amino acids were added via the tools available in Swiss-PdbViewer[11]. The resulting structure model was the basis for all modeling of the apo-state of mGBP2 ($mGBP2_{apo}$).

**GTP-bound model of mGBP2.** To obtain mGBP2 with GTP bound to it, the following steps were conducted. First, we had to provide the GTP parameters for the Amber99SB*-ILNDP force field in GROMACS format, as GROMACS was used for realizing the MD simulations. The original GTP parameters apart from the atomic charges were taken from the work of Meagher et al.[12] and incorporated into GROMACS following the procedure by Gao and Eriksson[13]. The partial atomic charges of GTP had to be determined to agree with the Amber99SB*-ILNDP charge assignment philosophy. For this, we followed the procedure that was used for the parametrization of the Amber ff99SB-ILDN force field, which employed the restrained electrostatic potential (RESP) method[14,15] based on a Hartree-Fock calculation, using a 6-31G* basis set, of geometry-optimized GTP. The geometry optimization was realized with a density functional-theory calculation with a B3LYP functional and a 6-31G* basis set. These quantum chemical calculations were performed using the Gaussian 09 program[16], following the recommendations of the Department of Theoretical Chemistry of the Lund University (available at `http://www.teokem.lu.se/ulf/Methods/resp.html`) and being in line with procedures described elsewhere[17,18]. The complete parameter file for use in GROMACS was built by combining the derived RESP charges and the GTP parameters available in AMBER frcmod file format[12] via the antechamber[19,20] and tleap[21] tools available in the AmberTools15 software package. The resulting prmtop and prmcrd files were converted to GROMACS topology and coordinate files using the ACPYPE (Ante Chamber PYthon Parser interface) tool[22].

To obtain a protein model of GTP-bound mGBP2, we made use of the crystal structure of the G domain of hGBP1 co-crystallized with the GTP-analogue GppNHp (PDB ID 2BC9)[23]. For homology modeling, the same procedure as explained above was employed, resulting in a homology model of the G domain of mGBP2, to which the M and E domain of the $mGBP2_{apo}$ model had to be added. Here, care had to be taken to avoid clashes between the loop involving residues 155–170 and the $\alpha$12-helix. This was accomplished by slightly stretching the bond between the C and $C_\alpha$-atoms of residue 481, which was enough to move the $\alpha$12/13-helices sufficiently away from the G domain. All these transformations were done using the VMD software[24]. Finally, the nucleotide GTP and the cation $Mg^{2+}$ as co-factor had to be added. To do so, the model was further processed with the free version of the Maestro program[25]. The structure model was loaded together with the superposed coordinates of $Mg^{2+}$ and GDPxAlF3 as available in the crystal structure of the G domain of hGBP1 co-crystalized with GDP-$AlF_3$ (PDB ID 2B92)[23]. To convert GDP-$AlF_3$ to GTP, $AlF_3$ was removed and a phosphate group was attached to the $\beta$-phosphate of the existing GDP. The $\gamma$-phosphate was added in a way as to avoid atom clashes and be in a reasonable position relative to important protein residues, in particular to K51 and the $Mg^{2+}$ co-factor, as described by Kravets[5]. The GTP structure was protonated to mimic the physiological pH value of 7.4, making use of the protonation state of GTP at pH 7.0 as available in the NMR-determined PDB structure with code 2KSQ, giving GTP an overall charge of $-4$. It was then pre-optimized according to the protein and co-factor environment within Maestro, which makes use of the OPLS 2005 force field[26]. The obtained coordinates defined the input structure for the simulations of GTP-bound mGBP2, denoted as $mGBP2_{GTP}$ in the paper.

**mGBP2 model involving the geranylgeranyl lipid anchor.** Similar to GTP, the force field parameters of the geranylgeranyl group together with C586, i.e., the residue to which it being attached, were generated using the Antechamber program[19,20] of the AmberTools15 software package[27] and ACPYPE[22]. The electron density calculations were performed using Gaussian 09[16] with the basis set 6-31G* at the Hartree-Fock (HF) level of theory. Partial charges were derived using the RESP method[14,15] following geometry optimization and the electrostatic potential calculations at the HF/6-31G* level. The resulting model involving both GTP in the G domain and the geranylgeranyl lipid anchor at the C-terminus represent the holo-state of mGBP2, thus denoted as mGBP2$_{\mathrm{holo}}$.

**mGBP2 dimer model.** To build the mGBP2$_{\mathrm{holo}}$ dimer, we superimposed two mGBP2$_{\mathrm{holo}}$ monomers on an energy-minimized structure of the hGBP1 dimer created by Barz et al.[28] using PyMOL[29]. This dimer structure was first equilibrated for 100 ns to test its stability and was found to be stable.

**mGBP2 monomer on a membrane.** For modeling membrane-bound mGBP2, we started by building a symmetric POPC (1-palmitoyl-2-oleoyl-*sn*-glycero-3-phosphocholine) membrane involving 1,418 lipids; this was accomplished with the CHARMM-GUI webserver[30,31]. To determine the influence of GTP on membrane-bound mGBP2, we created two systems: mGBP2 as mGBP2$_{\mathrm{holo}}$ and without GTP, both being placed on the membrane (denoted as mGBP2$^{\mathrm{GTP}}_{\mathrm{mem}}$ and mGBP2$^{\mathrm{noGTP}}_{\mathrm{mem}}$, respectively). The geranylgeranyl lipid anchor was inserted into the lipid bilayer by replacing one POPC molecule with the geranylgeranyl group, followed by a short energy minimization in vacuum to remove atom clashes. The parameters for POPC were taken from the Slipids force field[32,33].

To model a more realistic membrane, we employed CHARMM-GUI to create a symmetric membrane composed of three lipid types, namely 75% 1,2-dioleoyl-*sn*-glycero-3-phosphocholine (DOPC), 20% cholesterol, and 5% cardiolipin (CL), using experimental data as input (see below). The GTP-containing mGBP2 was placed on the membrane and anchored to it by inserting its geranylgeranyl group into the membrane, following the same protocol as for the POPC membrane systems. This system is denoted as mGBP2$^{\mathrm{GTP}}_{\mathrm{CL\text{-}mem}}$, and it was simulated using the CHARCHARMM36 force field[34,35]. The parameters for the geranylgeranyl group were derived using the CgenFF[36,37] at `https://cgenff.umaryland.edu/`. The latest parameters for the protein, GTP, and lipids were downloaded from the `http://mackerell.umaryland.edu/charmm_ff.shtml#gromacs` webserver.

#### MD simulation details

GROMACS 2016 or later versions[38,39] were employed for running all MD simulations.

**HREMD simulations of mGBP2 monomer in solution.** In order to identify the role of GTP and the geranylgeranyl group on the mGBP2 dynamics, we performed Hamilitonian replica exchange MD simulations for mGBP2$_{\mathrm{apo}}$, mGBP2$_{\mathrm{GTP}}$, and mGBP2$_{\mathrm{holo}}$. To this end, we placed each of the proteins in a rectangle simulation box of 8.5 nm × 9.0 nm× 17.0 nm and added ~40,000 water molecules as well as 11, 13, and 12 $Na^+$, respectively, for neutralization, resulting in around ~130,000 atoms. First, an energy minimization was performed using a steepest descent algorithm, followed by an $NpT$ (with $N$ as the number of atoms, $p = 1$ bar as the pressure, $T = 310$ K the temperature) position-restraint MD simulation of 500 ps in which the whole protein was restrained with a force constant of 10 kJ mol$^{-1}$ Å$^{-2}$ to equilibrate the solvent around the protein. Afterwards, the protein was slowly heated up to 310 K within a 500 ps MD simulation. Subsequently, a 20 ns $NpT$ equilibration was conducted, where we applied position restraints to the rigid $\beta$-sheets of the G domain to avoid overall protein rotation and translation. This allowed us to use a smaller simulation box and thus reduce the computational requirements, considering the substantial size of the protein and its elongated shape[28].

Next, we performed an HREMD simulation[40] with 16 replicas for each system. The energy function of mGBP2 including mGBP2–water interactions was modified in each replica by applying biasing factors of 310 K/$T$ with the 16 temperatures $T$ exponentially distributed between 310 and 370 K. This implies one unbiased replica, the so-called target replica at 310 K. This resulted in an average exchange probability between the neighbored replicas of ~50%. Each replica simulation was 200 ns long, leading to a total of 3.2 $\mu$s for the 16 replicas. To ensure that GTP stayed in its binding pocket, three distance restraints between GTP and K51, R48, and Y53 were applied using the pull code of GROMACS. For comparison, we also performed an HREMD simulation of mGBP2$_{\text{holo}}$ with 40 replicas involving a temperature range between 310 and 520 K and 400 ns per replica, amounting to 16 $\mu$s for that simulation. The HREMD simulations were conducted with Gromacs 2016.4 in combination with the PLUMED plugin (version 2.4.1 from `https://github.com/GiovanniBussi/plumed2/tree/v2.4`)[41].

In all HREMD simulations, we used the velocity rescaling thermostat and the isotropic Parrinello-Rahman barostat[42]. Electrostatic interactions were treated with the particle-mesh Ewald (PME) method[43,44] in conjunction with periodic boundary conditions and a real-space cutoff of 10 Å. The Lennard-Jones interactions were cut at 10 Å. The leapfrog stochastic dynamics integrator was used for the integration of equations of motion and the LINCS algorithm[45] for constraining all bond lengths. In the mGBP2$_{\text{apo}}$ system, virtual interaction sites were employed, permitting an integration time step of 4 fs while maintaining energy conservation[46]. For mGBP2$_{\text{GTP}}$ and mGBP2$_{\text{holo}}$, a time step of 2 fs was applied. Coordinates and velocities were saved every 10 ps.

**MD simulations of mGBP2$^{\text{holo}}$ as monomer and dimer in solution.** For the monomeric mGBP2$_{\text{holo}}$, we created the same system as for the HREMD simulation. The mGBP2$_{\text{holo}}$ dimer was solvated with 184,903 water molecules and the system neutralized with 24 $Na^+$, resulting in a total number of 573,673 atoms and a rectangular simulation box of 13.5 nm $\times$ 16.0 nm $\times$ 27.0 nm. The energy of both systems was first minimized using a steepest descent algorithm, followed by equilibration of the systems to the desired temperature of 310 K and pressure of 1 bar to mimic the physiological environment. First, a 0.1 ns $NVT$ equilibration was performed ($V$ being the box volume), followed by a 1 ns $NpT$ equilibration to adjust the pressure. During these steps, the protein's heavy atoms were restrained with a force constant of 10 kJ mol$^{-1}$ Å$^{-2}$, allowing the water molecules to relax around the solute. The velocity rescaling thermostat was employed to regulate the temperature in the $NVT$ simulations, while the Nosé-Hoover thermostat[47,48] and the isotropic Parrinello-Rahman barostat[42] were used for the $NpT$ simulations. The PME method[43,44] was used to calculate the electrostatic interactions with periodic boundary conditions. The cutoff for the van der Waals interactions and Coulombic interaction calculated in real space were set to 12 Å. The LINCS algorithm[45] was used to constrain all bond-lengths during the simulations. Production MD runs were performed for 1 $\mu$s for each system with position restraints on the rigid $\beta$-sheets of the G domain. The time step for integration was set to 2 fs, and the coordinates and velocities were saved every 20 ps.

**MD simulations of membrane-inserted mGBP2.** For the mGBP2$^{\text{GTP}}_{\text{mem}}$ and mGBP2$^{\text{noGTP}}_{\text{mem}}$ systems, we solvated the POPC membrane containing the geranylgeranyl-anchored protein, added 0.1 M NaCl as well as additional $Na^+$ to neutralize the overall charge of the system. The resulting box of 21 nm $\times$ 21 nm $\times$ 15 nm contains mGBP2$^{\text{GTP}}_{\text{mem}}$/mGBP2$^{\text{noGTP}}_{\text{mem}}$ as well as 1,417 POPC lipids, 437/435 $Na^+$ and 425 $Cl^-$ ions, and water, resulting in a total of 662,603/659,946 particles. To minimize the energy, equilibrate the systems and perform the 1 $\mu$s production runs, the same protocol as for the MD simulations of the mGBP2$_{\text{holo}}$ monomer and dimer was used, with the exception that a semi-isotropic Parrinello-Rahman barostat[42] was employed which is common for membrane systems. Moreover, no positional restraints were applied to the G domain, as the protein is already anchored. The same MD protocol was applied to the mGBP2$^{\text{GTP}}_{\text{CL-mem}}$ system. It involved a simulation box of size 19.8 nm $\times$ 19.8 nm $\times$ 17 nm containing 1,020 DOPC, 272 cholesterol and 68 cardiolipin lipids, 158,802 water molecules, 549 $Na^+$ and 401 $Cl^-$ ions, which is equivalent to 664,034 particles including the protein.

#### Experimental methods

**GFP-Trap®.** The one-step immunoprecipitation of GFP-tagged proteins is performed by anti-GFP VHH coupled agarose beads (90 $\mu$m) (Chromotek) with a binding capacity of 3-4 $\mu$g/10 $\mu$l GFP-Trap® slurry. For one immunoprecipitation reaction 106 MEF cells expressing GFP-tagged mGBP2 is recommended. Cell pellet was resuspended in 200 $\mu$l ice-cold RIPA buffer (10 mM Tris/Cl pH 7.5; 150 mM NaCl; 0,5 mM EDTA; 0,1% SDS; 1% Triton X-100; 1% Deoxycholate) supplemented with 1 mg/ml DNase, 2.5 mM MgCl2, protease inhibitors and 1 mM PMSF. Tube was placed on ice for 30 min and pipetted every 10 min. Cell lysate was centrifuged at 13,000 g at 4 °C for 15 min and transferred to a precooled tube containing a spin column

(Chromotek) for a clean elution of the bound proteins. 300 $\mu$l dilution buffer (10 mM Tris/Cl pH 7.5; 150 mM NaCl; 0.5 mM EDTA with 1 mM PMSF and protease inhibitors) and 25 $\mu$l GFP bead slurry (pre-washed and equilibrated in ice-cold dilution buffer) were added to the cell lysate in a spin column and incubated in end-over-end mixing for 2 h at 4 °C. Afterwards, tubes were centrifuged at 3,000 g for 2 min at 4v and 50 $\mu$l supernatant was saved for immunoblot analysis (non-bound sample). Beads with immunoprecipitated proteins were washed in ice-cold dilution buffer and 50 $\mu$l 0.2 M glycine elution buffer pH 2.5 was added and incubated for 30 sec under constant mixing. The spin column transferred to a new tube containing 5 $\mu$l 1 M Tris base pH 10.4 for neutralization, was centrifuged at 3,000 g for 3 min at 4 °C. Protein concentration of the supernatants including the eluted proteins of interest were measured using BCA protein assay kit (Pierce, Rockford, IL) and analyzed and verified via Western blot analyses and Coomassie blue stained SDS-PAGE.

**Giant unilammelar vesicle preparation** Giant unilammelar vesicles (GUVs) were prepared by electroformation (11Hz, 2V for 3 h at room temperature) using a construction scaffold of 74.75 mol% DOPC (1,2-Dioleoyl-sn-glycero-3-phosphocholine), 20 mol% cholesterol, 0.25 mol% TexasRed and 5 mol% lipid of interest (cardiolipin solved in chloroform). Lipid extracts were used from Avanti Polar Lipids. For each GUV preparation, two ITO-coated glass slides (Glas & Optik GmbH, Iserlohn, Germany, #CEC010S) were cleaned two times by EtOH followed by two times with chloroform. Within a circular area on the ITO-coated glass side the lipid solution was spaced out evenly in a total volume of 20 $\mu$l of each slide using a Hamilton™ syringe. For the rehydration of lipid films, the circular lipid area on each slide is fenced by a Vitrex™ ring. The lipid coated sides of each slide were faced to each other and pressed on the Vitrex™ ring. Inside of this Vitrex™ chamber, 400 $\mu$l of 10% sucrose solution are injected. After the electroformation for 3 h, the sucrose buffer containing GUVs were harvested and transferred into a 2 ml Eppendorf tube. The GUV solution contains vesicles with different diameters of 5–60 $\mu$m.

#### Supplementary figures

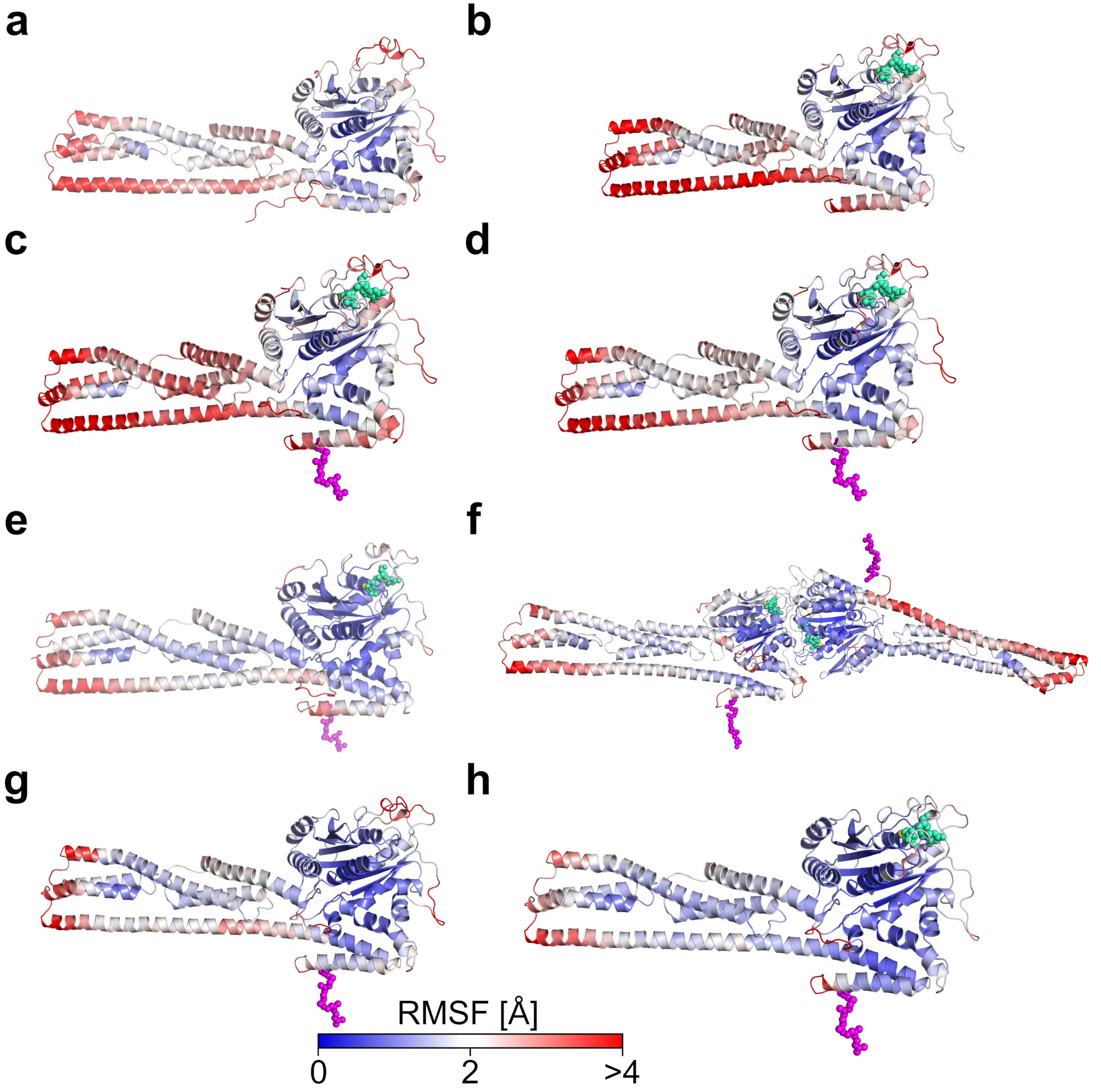


**Supplementary Figure S1. Structural flexibility of the different mGBP2 systems.** The flexibility, quantified as the RMSF of the $C_\alpha$ atom during the simulations, is projected onto the mGBP2 starting structures for **(a)** mGBP2$_{apo}$, **(b)** mGBP2$_{GTP}$, and **(c)** mGBP2$_{holo}$ for the 16 $\times$ 200 ns HREMD, **(d)** mGBP2$_{holo}$ for the 40 $\times$ 400 ns HREMD, **(e)** mGBP2$_{holo}$ monomer and **(f)** mGBP2$_{holo}$ dimer for 1 $\mu$s MD, as well as **(g)** mGBP2$^{noGTP}_{mem}$ and **(h)** mGBP2$^{GTP}_{mem}$ monomer on the membrane for 1 $\mu$s MD. Rigid residues are depicted in blue, whereas the flexible residues are shown in red, according to the color scale at the bottom.

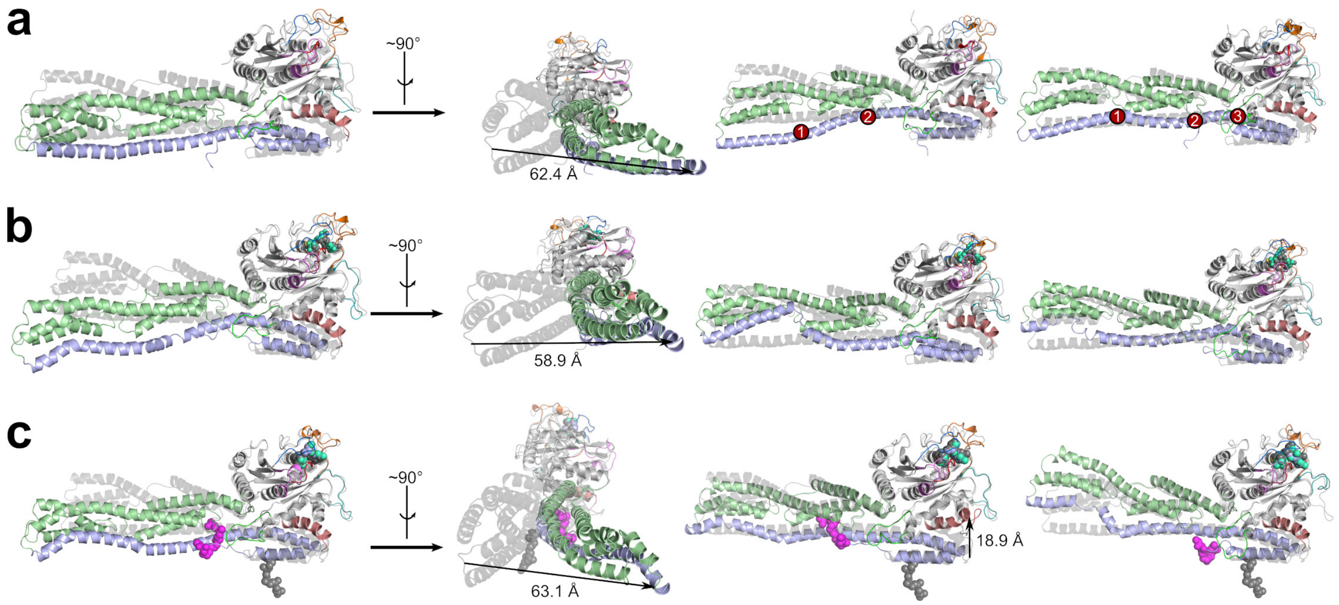


**Supplementary Figure S2. Movements and structural changes of M/E domain and $\alpha$4'-helix in the 16 $\times$ 200 ns HREMD simulations.** The conformations with the largest deviations from the start structure are shown for mGBP2$_{apo}$ in **(a)**, mGBP2$_{GTP}$ in **(b)** and mGBP2$_{holo}$ in **(c)**, using residue L480 for measuring the motions of the M/E domain and K212 for assessing the motions of $\alpha$4' of the G domain. Preferred places of kinks and temporary helix unfolding in $\alpha$12 are highlighted by red circles in **(b)**.

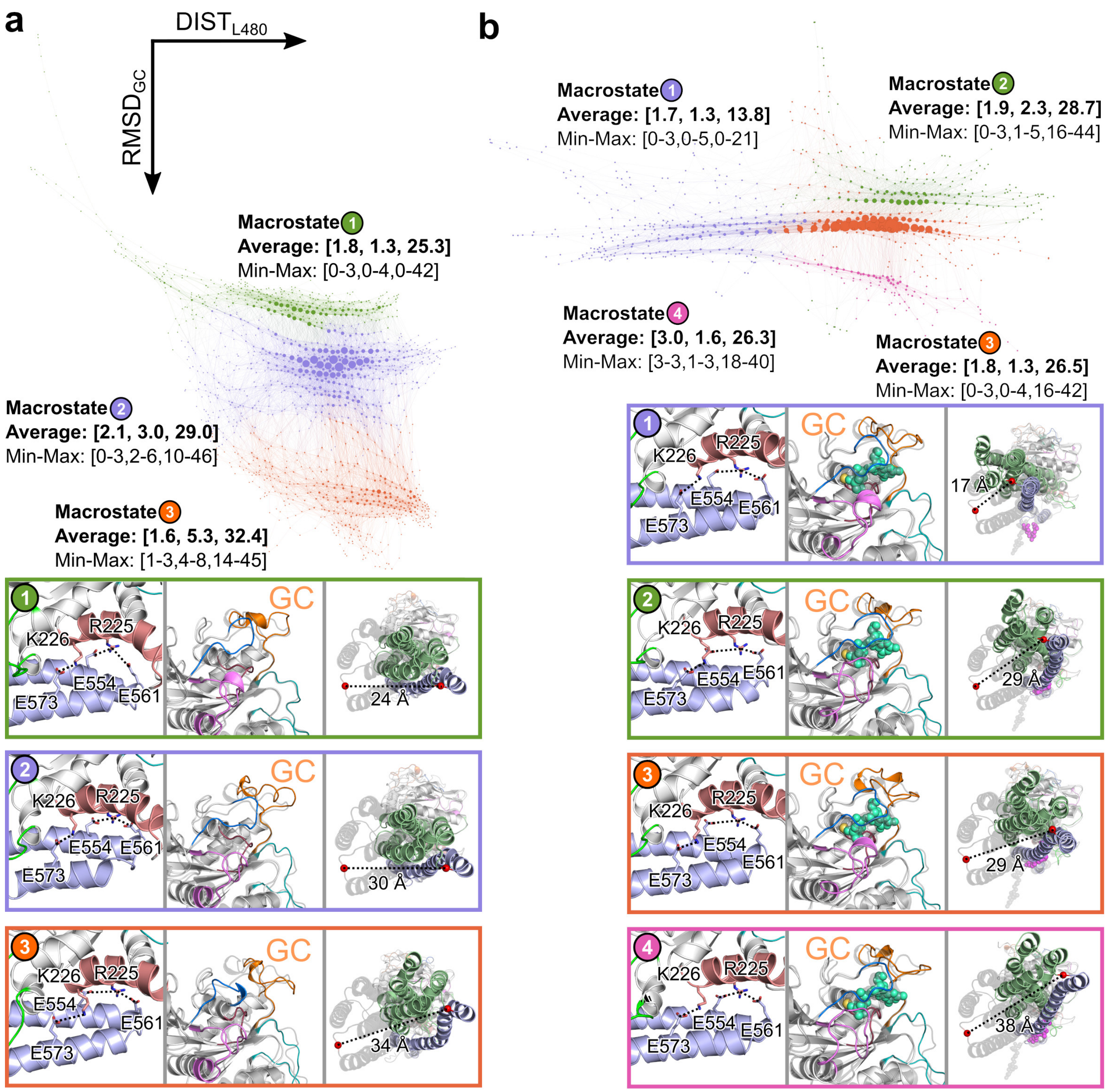


**Supplementary Figure S3. Transition networks of (a) mGBP2$_{apo}$ and (b) and mGBP2$_{holo}$.** The TNs were calculated using three descriptors: (i) the presence of three salt bridges (SB) between the G and E domain (R225–E554, R225–E561, K226–E573); (ii) the RMSD of the guanine cap (RMSD$_{GC}$); (iii) the distance of the C$_\alpha$ atom of L480 between the MD and the start structure ($d_{480}$). The states of the TNs are arranged such that major $d_{480}$ changes occur along the $x$-axis and major RMSD$_{GC}$ changes along the $y$-axis. The size of the nodes reflect the population of the corresponding state. The networks are further divided into macrostates, which were determined using the modularity class feature of Gephi that identifies local communities of highly interconnected states. The average and minimum/maximum values of the descriptors per macrostate are provided as [SB, RMSD$_{GC}$, $d_{480}$]. In addition, for each macrostate, a representative conformation was extracted and its structural elements underlying descriptor definitions (i) to (iii) (from left to right) are shown along with the starting structure (for RMSD$_{GC}$ and $d_{480}$, in gray) to illustrate the movement in question. The salt bridges and guanine cap are labeled, the C$_\alpha$ atom of L480 is shown as red sphere, and the $d_{480}$ value is given.

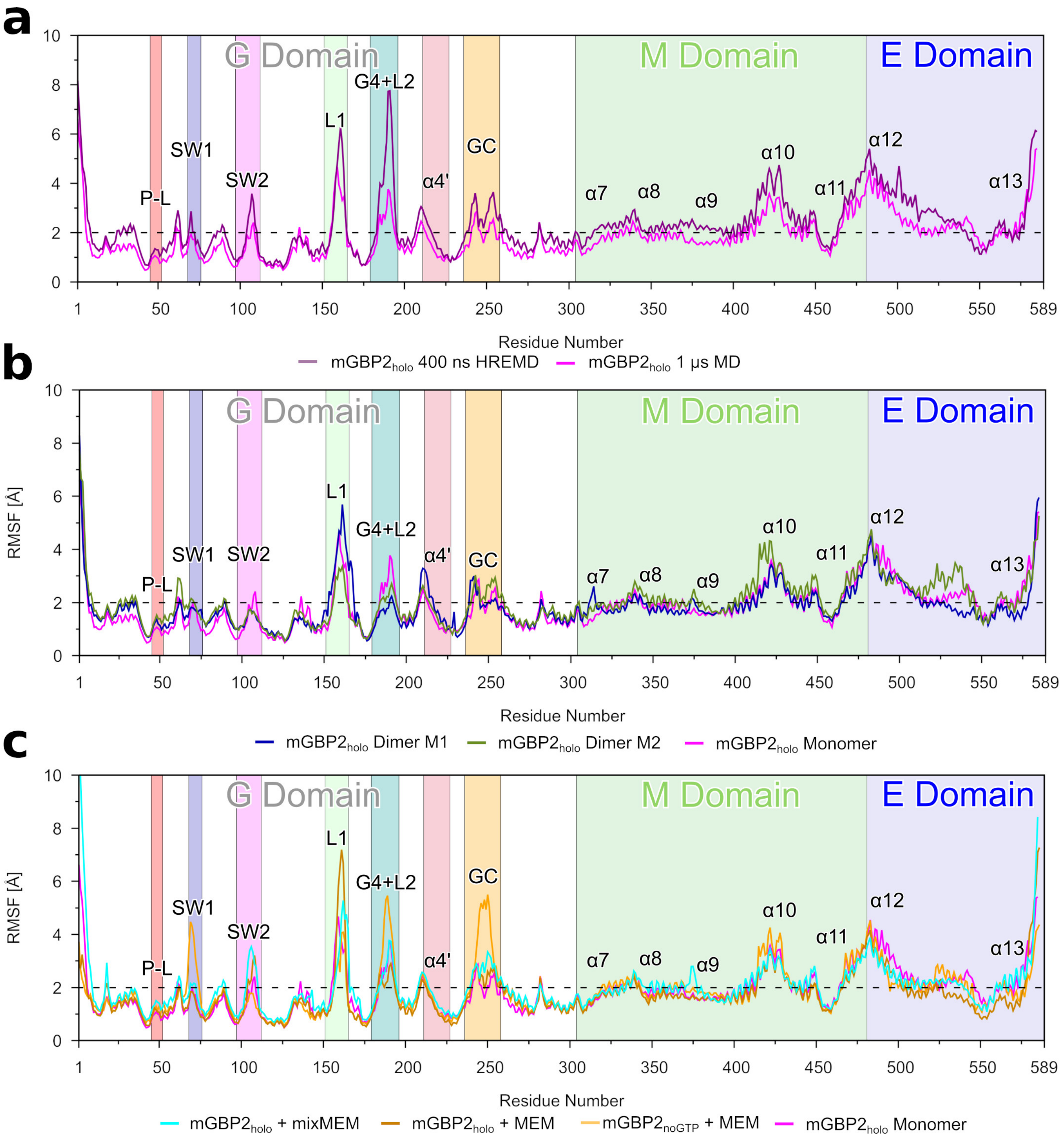


**Supplementary Figure S4. The root mean square fluctuations per residue of mGBP2 in different states during 40 × 400 ns HREMD simulations and 1 $\mu$ MD simulations.** **(a)** The fluctuations of mGBP2$_{\mathrm{holo}}$ in the target replica of the HREMD simulation (dark violet) and the MD simulation (magenta). The latter is repeated in panel **(b)** and serves as reference for the fluctuations of mGBP2$_{\mathrm{holo}}$ as part of the dimer (monomer M1: blue; monomer M2: green) as well as in **(c)** where it is compared with mGBP2$^{\mathrm{GTP}}_{\mathrm{CL\text{-}mem}}$ (cyan), mGBP2$^{\mathrm{GTP}}_{\mathrm{mem}}$ (brown) and mGBP2$^{\mathrm{noGTP}}_{\mathrm{mem}}$ (orange). All important motifs, loops and helices are labeled and a background color added using the same colors as for the corresponding structural units as in Fig. 1.

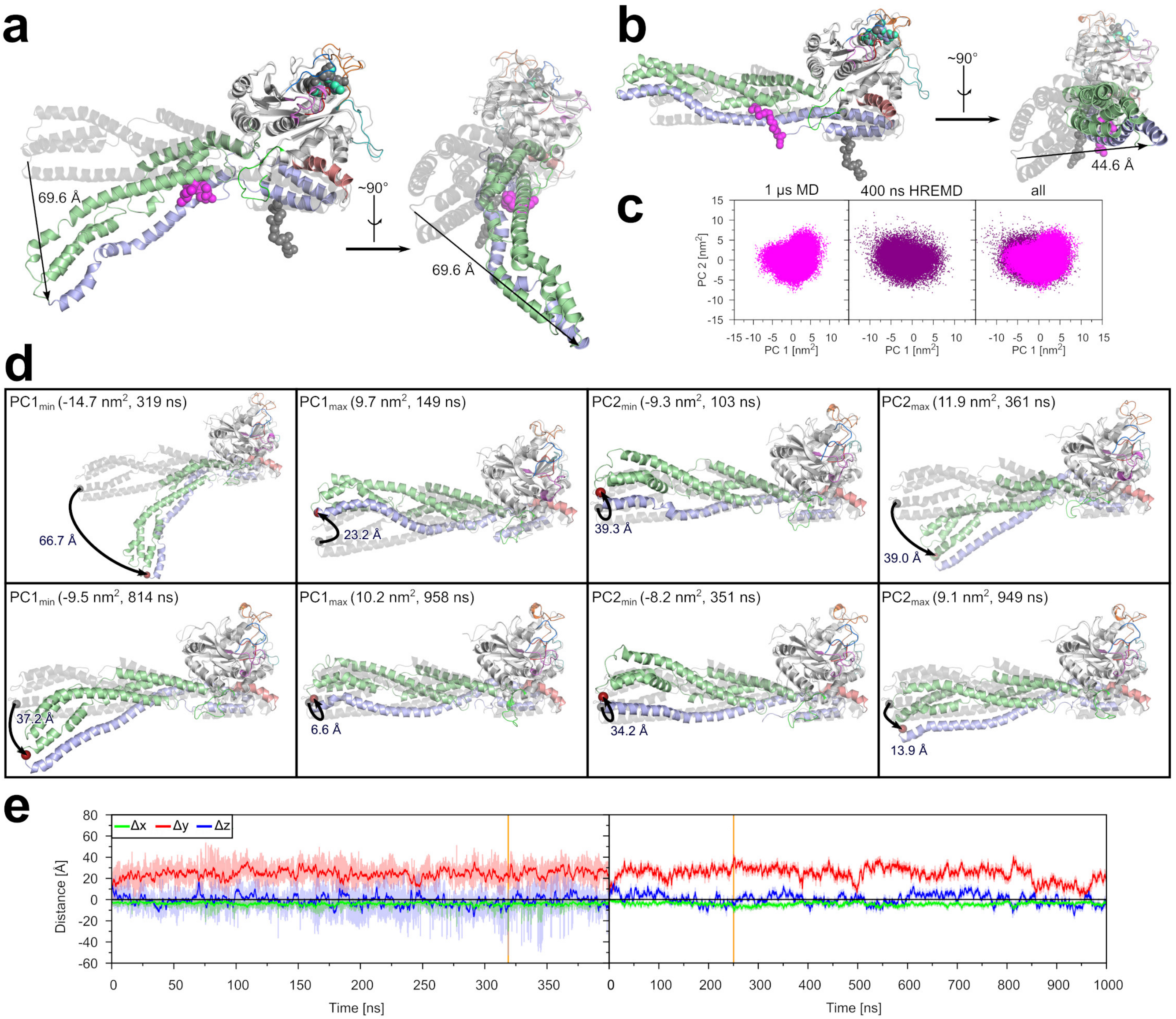


**Supplementary Figure S5. Movement and structural changes of the M/E domain during the 40 × 400 ns HREMD simulation and 1 $\mu$s MD of the mGBP2$_{\mathrm{holo}}$ monomer.** The conformation with the largest deviation from the start structure, as measured by the motions of L480 at the tip of the M/E domain, is shown for the HREMD simulation in **(a)** and the MD simulation in **(b)**. **(c)** The projection of the target replica of the HREMD simulation (violet) and of the MD simulation (magenta) along the first two principal components describing the motions of the M/E domain. **(d)** Structures corresponding to the minimal and maximal PC1 and PC2 values extracted from the HREMD simulation (upper panel) and the MD simulation (lower panel) together with the starting structure of the simulations serving as reference. The deviations of L480 (red sphere) from the starting structure are indicated. **(e)** The motions of L480 are further assessed by monitoring its coordinate distances $\Delta x$ (green), $\Delta y$ (red) and $\Delta z$ (blue) from the reference structure in the target replica of the HREMD simulation (left) and in the MD simulation (right). The solid lines show the running average and the shades illustrate the raw data.

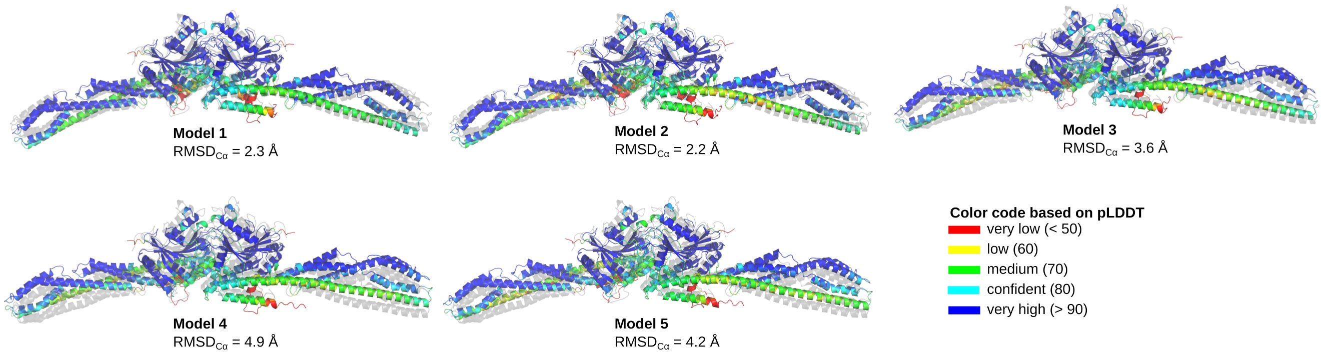


**Supplementary Figure S6. The five most likely mGBP2 dimer models as predicted by AlphaFold-Multimer.** The dimer models are colored based on a per-residue estimate of the prediction's confidence (called pLDDT) on a scale from 0–100. Regions with pLDDT $> 90$ are shown in blue and expected to be modeled to high accuracy. Regions with pLDDT between 70 and 90 are expected to be modeled well (a generally good backbone prediction), while the predictions for regions with pLDDT between 50 and 70 are of low confidence and should be treated with caution. The 3D coordinates of regions with pLDDT $< 50$ should not be interpreted. Such low pLDDT values are a strong predictor of disorder or that the region in question is only structured as part of a complex. The G and M domains in the AlphaFold-Multimer models are predicted with high confidence, while it is lower for the E domain. The deviation from the mGBP2 dimer that we constructed based on the X-ray structure of the G-domain dimer of hGBP1 (shown as gray cartoon) is provided in terms of the RMSD of the $C_\alpha$ atoms.

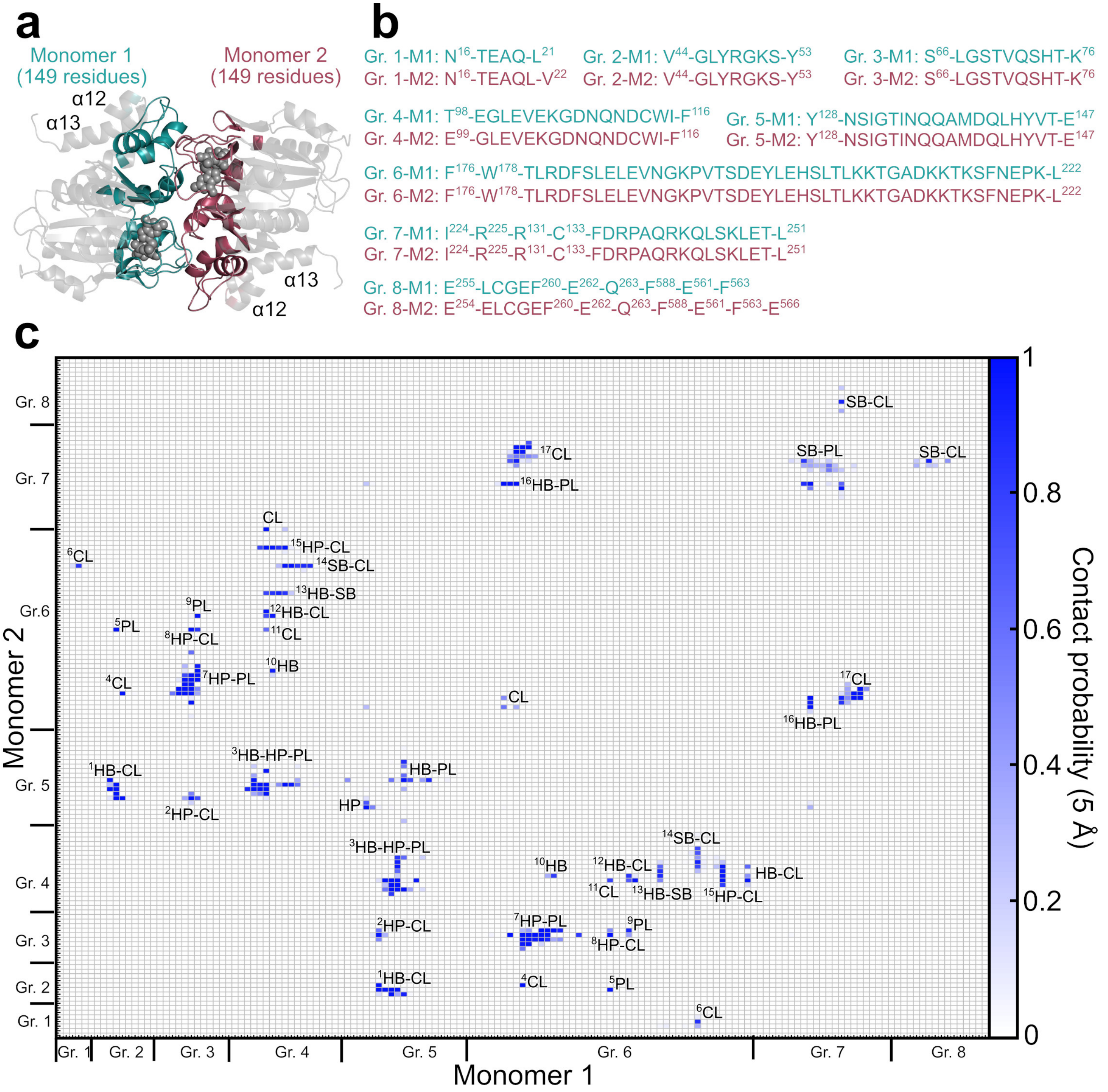


**Supplementary Figure S7. Interaction interface of the mGBP2$_{holo}$ dimer.** **(a)** All residues within 12 Å of the dimer interface are highlighted in dark teal for monomer M1 of the dimer and dark red for monomer M2 of the dimer. These residues are mainly from the G domain and $\alpha$12/13 of the E domain. GTP is shown as gray spheres, while the parts of the G domain and $\alpha$12/13 that are not involved in the protein–protein interactions are represented as transparent gray cartoon. **(b)** List of the residues involved in the interactions. They are combined into eight groups, which are used as labels in **(c)** where the probabilities of residue–residue contacts between M1 and M2 are shown. The probabilities were calculated considering contacts with a maximum distance of 5 Å between the residues. They are shown as a heat map, in which the depth of the blue color represents the height of the contact probability between 0 and 1. The nature of the contacts is provided, distinguishing between hydrogen bonds (HB), salt bridges (SB), interactions between hydrophobic residues (HP), interactions between polar residues (PL), $\pi$–$\pi$ interactions (PP), and residues that are close to each other but do not belong to any of the special interactions (CL). The consecutive superscripts indicate symmetric interactions.

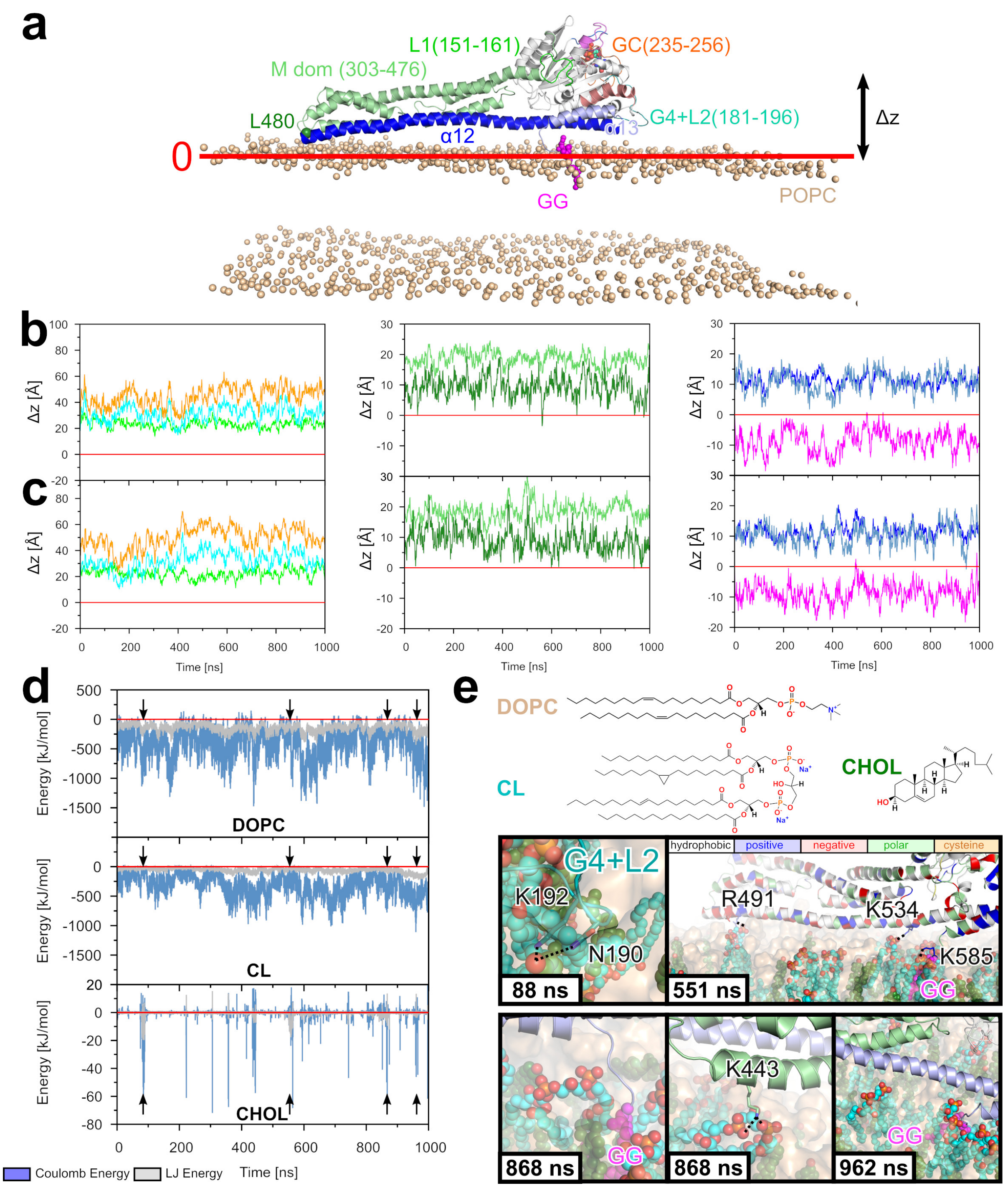


**Supplementary Figure S8. Motions and interactions of membrane-bound mGBP2.** **(a)** The motions are measured by distances along the *z*-coordinate ($\Delta z$) between the centers of mass of selected residue groups and the average position of the lipid head groups, as indicated by the red line (defining $z = 0$). In **(b)** for mGBP2$^{\text{noGTP}}_{\text{mem}}$ and in **(c)** for mGBP2$^{\text{GTP}}_{\text{mem}}$, the evolution of $\Delta z$ for the different residue groups relative to the POPC membrane are shown. The colors are the same as those used in **(a)** for defining the residue groups. The results are separately shown for the residue groups of the different domains (left: G domain; middle: M domain; right: E domain and geranylgeranyl (GG) anchor). **(d)** The evolution of the interaction energies between mGBP2 and the different lipids of the DOPC/cholesterol/cardiolipin membrane, decomposed into $E_{\text{Coul}}$ (blue) and $E_{\text{LJ}}$ (gray), is shown. The arrows indicate the times at which the structures displayed in **(e)** were extracted: 88, 551, 868, and 962 ns. The 2D structures of the lipids are provided above these MD snaphots taken from the mGBP2$^{\text{GTP}}_{\text{CL-mem}}$ simulation. In the structural plots on the bottom, DOPC is shown in ochre, cholesterol (CHOL) in green, and cardiolipin (CL) in cyan. important residues in interaction with the membrane as well as the geranylgeranyl (GG) anchor are labeled.

#### Supplementary tables

**Supplementary Table S1. Structural elements and function of the important motifs, loops and helices for all three domains of mGBP2.** Abbreviations: P-L = phosphate-binding loop, SW1/2 = switch 1/2, L1/2 = loop 1/2, G4 = G4 motif X(V/L)RD, GC = guanine cap, GG = geranylgeranyl.

| Motif | Sequence | Key residues | Function |
|---|---|---|---|
| | | **G domain (residues 1–303)** | |
| P-L | 44–52 | R48, K51 | GTP binding and hydrolysis (internal arginine finger) |
| SW1 | 68–76 | S73, T75 | GTP binding and hydrolysis ($Mg^{2+}$ and $\gamma$-phosphate) |
| SW2 | 97–112 | E99 | GTP binding and hydrolysis |
| L1 | 151–165 | | |
| G4+L2 | 181–196 | D182 | GTP binding and specificity |
| $\alpha$4’ | 211–227 | R225, K226 | Contact between G and E domain |
| GC | 235–256 | | Dimerization interface of mGBP2 |
| | | **M domain (residues 304–481)** | |
| $\alpha$7 | 310–340 | | Helix bundle 1 |
| $\alpha$8 | 348–369 | | Helix bundle 1 |
| $\alpha$9 | 377–422 | | Long helix as connector of the two helix bundles |
| $\alpha$10 | 430–447 | | Helix bundle 2 |
| $\alpha$11 | 455–476 | | Helix bundle 2 |
| | | **E domain (residues 482–589)** | |
| $\alpha$12 | 482–561 | E554, E561 | Contact with membrane and helix $\alpha$4’ of G domain |
| $\alpha$13 | 565–585 | E566, E573 | Contact with membrane and helix $\alpha$4’ of G domain |
| GG | 586–589 | C586 | Geranylgeranyl lipid anchor with CaaX (CTIL) motif |

**Supplementary Table S2. Summary of different properties of the monomeric and dimeric systems.**

| Properties | $mGBP2_{apo}$ | $mGBP2_{GTP}$ | $mGBP2_{holo}$ | $mGBP2_{holo}$ | $mGBP2_{holo}$ | $mGBP2_{holo}$-dimer[a] |
|---|---|---|---|---|---|---|
| | 16 × 200 ns HREMD | | | 40 × 400 ns HREMD | 1 $\mu$s MD | 1 $\mu$s MD |
| PC1 [$nm^2$] | −12.1 to 8.5 | −12.2 to 12.7 | −12.1 to 16.9 | −14.7 to 9.7 | −9.5 to 10.2 | — |
| PC2 [$nm^2$] | −8.1 to 10.9 | −8.9 to 10.6 | −9.0 to 9.4 | −9.3 to 11.9 | −8.2 to 9.1 | — |
| $\Delta x$ [Å][b] | −4.2 ± 0.02 (−21.2 to 2.2) | −3.9 ± 0.02 (−18.3 to 2.4) | −3.8 ± 0.02 (−19.6 to 3.2) | −3.9 ± 0.02 (−30.1 to 2.4) | −4.2 ± 0.01 (−13.2 to 1.3) | −14.9 ± 0.02 & 5.2 ± 0.01 (−35.8 to 3.0 & −2.0 to 16.2) |
| $\Delta y$ [Å][b] | 26.0 ± 0.07 (−8.1 to 55.6) | 23.3 ± 0.07 (−19.0 to 52.9) | 22.2 ± 0.07 (−13.6 to 56.3) | 24.2 ± 0.05 (−5.6 to 54.0) | 24.6 ± 0.03 (−1.8 to 42.9) | 15.5 ± 0.03 & −6.8 ± 0.03 (−9.3 to 36.3 & −37.4 to 18.5) |
| $\Delta z$ [Å][b] | −6.4 ± 0.05 (−32.2 to 19.4) | −2.3 ± 0.06 (−34.8 to 25.3) | −0.59 ± 0.09 (−37.7 to 42.1) | −0.99 ± 0.07 (−52.8 to 27.2) | 0.37 ± 0.02 (−18.5 to 18.5) | −22.5 ± 0.02 & 12.7 ± 0.04 (−38.5 to 4.7 & −17.6 to 36.1) |
| L480–L480 [Å][b] | 28.2 ± 0.07 (1.2 to 62.4) | 25.3 ± 0.07 (0.7 to 58.9) | 25.9 ± 0.06 (1.2 to 63.1) | 26.5 ± 0.05 (1.9 to 69.6) | 25.6 ± 0.03 (0.6 to 44.6) | 31.8 ± 0.03 & 17.6 ± 0.03 (0.0 to 54.5 & 0.0 to 41.3) |
| $\alpha_E$[c] [%] | 88.0 | 84.1 | 85.1 | 84.1 | 86.1 | 87.0 & 84.1 |

[a] The values are given for monomer 1 and 2.
[c] The mean ± standard error of the mean as well as the minimal to maximal values in brackets below are listed.
[d] $\alpha_E$ denotes the time-averaged helix content of the E domain.

**Supplementary Table S3. Flexibility of the loops of the G domain, helix $\alpha$4' and the geranylgeranyl anchor determined by clustering of the conformations of the respective 1 $\mu$s MD simulations or the target replica of the 40 $\times$ 400 ns HREMD simulation.**

| | $mGBP2_{holo}$ monomer, 1 $\mu$s MD | | | $mGBP2_{holo}$ monomer, 400 ns HREMD | | |
|---|---|---|---|---|---|---|
| (Motif-)Loop | clusters | population[%][a] | RMSD[Å][b] | clusters | population[%][a] | RMSD[Å][b] |
| P-L | 1 | 100 | – | 1 | 100 | – |
| SW1 | 1 | 100 | – | 2 | 100 | 3.1 |
| SW2 | 3 | 100 | 5.0 | 3 | 100 | 4.9 |
| L1 | 68 | 81.3 | 13.5 | 39 | 76.5 | 13.3 |
| G4+L2 | 2 | 100 | 2.8 | 4 | 99.9 | 4.3 |
| $\alpha$4' | 2 | 100 | 4.0 | 3 | 100 | 5.7 |
| GC | 6 | 99.8 | 5.2 | 6 | 99.6 | 5.1 |
| GG | 59 | 67.4 | 6.6 | 97 | 32 | 6.5 |

| | M1 of the $mGBP2_{holo}$ dimer, 1 $\mu$s MD | | | M2 of the $mGBP2_{holo}$ dimer, 1 $\mu$s MD | | |
|---|---|---|---|---|---|---|
| (Motif-)Loop | clusters | population[%][a] | RMSD[Å][b] | clusters | population[%][a] | RMSD[Å][b] |
| P-L | 1 | 100 | – | 1 | 100 | – |
| SW1 | 1 | 100 | – | 1 | 100 | – |
| SW2 | 1 | 100 | – | 1 | 100 | – |
| L1 | 34 | 72.0 | 13.2 | 15 | 95.0 | 12.8 |
| G4+L2 | 1 | 100 | – | 1 | 100 | – |
| $\alpha$4' | 3 | 100 | 3.6 | 2 | 100 | 3.9 |
| GC | 2 | 100 | 2.2 | 3 | 100 | 4.9 |
| GG | 72 | 48.7 | 6.7 | 106 | 31.3 | 6.5 |

| | $mGBP2^{noGTP}_{mem}$, 1 $\mu$s MD | | | $mGBP2^{GTP}_{mem}$, 1 $\mu$s MD | | |
|---|---|---|---|---|---|---|
| (Motif-)Loop | clusters | population[%][a] | RMSD[Å][b] | clusters | population[%][a] | RMSD[Å][b] |
| P-L | 1 | 100 | – | 1 | 100 | – |
| SW1 | 34 | 67.2 | 12.8 | 4 | 99.9 | 6.2 |
| SW2 | 1 | 100 | – | 1 | 100 | – |
| L1 | 29 | 81.2 | 11.5 | 20 | 86.8 | 10.6 |
| G4+L2 | 62 | 60.6 | 12.9 | 11 | 97.6 | 9.4 |
| $\alpha$4' | 3 | 100 | 5.2 | 3 | 100 | 5.1 |
| GC | 101 | 44.4 | 19.6 | 5 | 99.9 | 5.1 |
| GG | 75 | 51.1 | 6.5 | 80 | 47.1 | 6.8 |

[a] Percentage of the structures which are cumulatively represented by the first three clusters.
[b] The largest RMSD found between the clusters.

**Supplementary Table S4. Interactions between the residues in the mGBP2$_{holo}$ dimer interface.**

| No.[a] | M1[b] | M2[b] | Interaction[c] | Stability[d] |
|---|---|---|---|---|
| 1* | Y47 (P-L) | I134 | HB-SC-BB | very stable |
| 2* | Y47 (P-L) | Y199 | PL-SC-SC | very stable |
| 3 | V71 (SW1) | T134 | HP | aliphatic groups, very stable |
| 4 | V71 (SW1) | Y199 | HP | stable |
| 5 | Q72 (SW1) | H202 | HB-SC-SC | stable |
| 6* | G68–Q72 (SW1) | F183–G191 (G4+L2) | HP-PL | stable |
| 7* | E102 (SW2) | Q137 | HB-SC-SC | unstable |
| 8* | E102 (SW2) | Q136 | HB-BB-BB | very stable |
| 9 | V104 (SW2) | I134/M139 | HP | very stable |
| 10 | V104 (SW2) | F217 ($\alpha$4’) | HP | stable |
| 11* | E105 (SW2) | N190/H202 (G4+L2) | HB-SC-SC | very stable; HB network |
| 12 | E105 (SW2) | K207 | SB-SC-SC | very unstable |
| 13* | K106 (SW2) | K207 | HB-BB-SC | unstable |
| 14* | D108 (SW2) | K213 ($\alpha$4’) | SB | stable |
| 15* | – | F217/K221 ($\alpha$4’) | PP | very stable cation-$\pi$-stacking |
| 16 | N135 | Q137 | HB-SC-SC | unstable |
| 17* | Q137 | Q137 | HB-SC-BB | unstable |
| 18* | D182 (G4+L2) | R238 (GC) | HB-BB-SC | stable |
| 19* | S184 (G4+L2) | R238 (GC) | PL-SC | stable |
| 20 | R221 ($\alpha$4’) | V104 (SW2) | HB-SC-BB | stable |
| 21 | D237 (GC) | R242 (GC) | SB | very unstable |
| 22* | D237 (GC) | K243 (GC) | SB | stable |
| 23 | R238 (GC) | D237 (GC) | SB | very unstable |
| 24 | K243 (GC) | E259 | SB | stable |
| 25 | E259 | R242/K243 (GC) | SB | stable, alternated |

[a] The *-sign indicates that the interactions appear between both monomers M1 to M2 and M2 to M1 which means they are symmetric.
[b] If the residue belongs to a specific motif or helix, then it is written in the brackets.
[c] The nature of the interactions is provided, distinguishing between hydrogen bonds (HB), salt bridges (SB), interactions between hydrophobic residues (HP), interactions between polar residues (PL), and $\pi$–$\pi$ interactions (PP). To further specify the interactions, we also used the difference between backbone (BB) and side chain (SC), where the order is "type of interaction-M1(BB or SC)-M2(BB or SC)".
[d] Very stable interactions are present the whole time (contact probability (CP) $\geq 0.9$), while stable interactions are present in at least half of the simulation time ($0.5 \leq$ CP $< 0.9$). Unstable interactions are in the range of $0.25 \leq$ CP $< 0.5$, whereas very unstable implies CP $< 0.25$. The probabilities were calculated considering contacts with a maximum distance of 5 Å between the residues.

**Supplementary Table S5. Average motions of residue L480 at the M/E tip with respect to the MD starting structure as well as motions of residue groups with respect to the membrane surface in 1 $\mu$s MD simulations of mGBP2$_{\text{mem}}^{\text{noGTP}}$ and mGBP2$_{\text{mem}}^{\text{GTP}}$.** The motions of L480 are provided as Cartesian displacements $\Delta x$, $\Delta y$ and $\Delta z$ as well as its distance from the starting structure. The motions with respect to the membrane surface are given by $\Delta z$ values of the centers of mass of the respective residue group, where $z$ is the direction of the membrane normal and $z = 0$ defines the membrane surface given by the lipid headgroups.

| **L480 motions [Å]** | **mGBP2$_{\text{mem}}^{\text{noGTP}}$** [a] | **mGBP2$_{\text{mem}}^{\text{GTP}}$** [a] |
|---|---|---|
| $\Delta x$ | $-4.0 \pm 0.01$ ($-15.0$ to 3.6) | $-3.8 \pm 0.01$ ($-17.0$ to 1.8) |
| $\Delta y$ | $21.7 \pm 0.05$ ($-16.9$ to 55.0) | $22.9 \pm 0.04$ ($-16.2$ to 56.7) |
| $\Delta z$ | $0.3 \pm 0.04$ ($-32.7$ to 33.9) | $1.7 \pm 0.04$ ($-28.3$ to 33.9) |
| distance | $24.7 \pm 0.04$ (0.8 to 57.0) | $25.0 \pm 0.04$ (0.8 to 59.1) |
| **$\Delta z$ motions of residue groups [Å]** | | |
| G domain: L1 (151–161) | $23.1 \pm 0.10$ (14.0 to 35.7) | $22.4 \pm 0.12$ (11.0 to 33.7) |
| G domain: G4+L2 (181–196) | $31.1 \pm 0.19$ (14.6 to 49.7) | $31.9 \pm 0.23$ (10.1 to 54.2) |
| G domain: GC (235–256) | $44.7 \pm 0.23$ (24.2 to 63.4) | $49.7 \pm 0.27$ (22.5 to 70.3) |
| M domain (303–476) | $18.6 \pm 0.07$ (11.5 to 25.1) | $18.7 \pm 0.09$ (9.2 to 30.1) |
| M/E tip (L480) | $9.3 \pm 0.12$ ($-3.6$ to 21.9) | $8.9 \pm 0.12$ ($-0.32$ to 24.0) |
| E domain: $\alpha$12 | $11.4 \pm 0.07$ (4.8 to 17.0) | $11.6 \pm 0.08$ (4.8 to 20.4) |
| E domain: $\alpha$13 | $10.8 \pm 0.11$ (1.7 to 19.9) | $10.3 \pm 0.11$ ($-0.95$ to 20.4) |
| E domain: geranylgeranyl anchor | $-8.6 \pm 0.12$ ($-18.6$ to 0.82) | $-8.8 \pm 0.09$ ($-18.4$ to 4.5) |

[a] The mean ± standard error of the mean as well as the minimal to maximal values in brackets below are listed.
